# Durable and reversible epigenetic programming of human hematopoietic stem cells as a platform for modulation of platelet function

**DOI:** 10.64898/2026.03.27.714536

**Authors:** Tianyi Ye, Wanying Xu, Maria N. Barrachina, Peng Lyu, Mateusz Antoszewski, Lucrezia della Volpe, Chun-Jie Guo, Andrew J. Lee, Emily Villalpando, Isaac A. Johnson, Madelaine S. Theardy, Spencer D. Shelton, Lara Wahlster, Alexis Caulier, Luana Messa, Michael Poeschla, Gaurav Agarwal, Ronodeep Mitra, Alec A. Schmaier, Jonathan S. Weissman, Kellie R. Machlus, Vijay G. Sankaran

**Author notes:** Correspondence should be addressed to V.G.S.

## Abstract

Programmable epigenome editing offers control of gene expression without altering DNA sequence, but its capacity to program human hematopoietic stem cells (HSCs) and their differentiated progeny remains incompletely characterized. Here, we establish that DNA methylation-based epigenome editors delivered transiently as RNA induce heritable gene silencing in primary human HSCs. Targeting the platelet integrin β3 (*ITGB3*) produced silencing that persisted through long-term HSC self-renewal and megakaryocytic differentiation, impaired aggregation of *in vitro*-derived platelets, and was partially reversed through targeted demethylation at this locus. Silencing remained stable in human HSCs four months after transplantation into mice. Targeting additional platelet-associated genes demonstrated broad applicability with this approach. Together, these findings establish DNA methylation-based epigenome editing as a platform for durable and reversible programming of human HSCs and demonstrate its application to modulating platelet function.

## Introduction

Hematopoietic stem cells (HSCs) reside within the bone marrow and are responsible for maintaining and replenishing the blood and immune systems throughout life.^1,2^ HSCs are unique, as they can be replaced through transplantation and therefore serve as an ideal target for one-time therapeutic interventions for a range of blood, immune, and metabolic disorders.^3^ Clinically, HSC transplantation has long been used to treat hematologic malignancies and inherited disorders, and recent advances in gene therapy and gene editing have further expanded its therapeutic reach.^3–6^ For example, the approval of Casgevy (exagamglogene autotemcel) for severe sickle cell disease and β-thalassemia marks the first clinical translation of targeted HSC engineering by genome editing, establishing a foundation for next-generation therapies.^7,8^ However, despite these advances, the therapeutic application of genome editing in HSCs has largely focused on select monogenic blood and immune disorders, and often applies tools that leave long-term scars in the genome, including undesired on and off-target edits, as well as other heritable gene expression alterations, which limit broader potential applications.^9–12^

Programmable epigenome silencers target specific genomic loci to modulate gene expression without altering the underlying DNA sequence, representing versatile and potentially safer therapeutic modalities to repress disease-linked genes. Leveraging epigenetic memory, the faithful maintenance of cytosine methylation at CpG sites through DNA replication and cell division, the initial hit-and-run targeted epigenetic silencing strategy and subsequent systems such as CRISPRoff and CHARM (Coupled Histone tail for Autoinhibition Release of Methyltransferase) establish sustained transcriptional repression in mammalian cells by depositing DNA methylation at targeted loci through exogenous DNMT3A catalytic domains or by recruiting endogenous DNMT3A together with DNMT3L, in addition to incorporating the KRAB repressor domain.^13–15^

These DNA methylation-based silencers have been shown to effectively and durably silence *PCSK9* in the liver to lower circulating cholesterol levels, the prion disease gene *Prnp* in neurons, specific genes to improve CAR-T persistence, as well as to achieve multiplexed silencing of clinically relevant genes in primary human T cells.^15–19^ Importantly, such silencing is reversible through targeted DNA demethylation, providing an additional layer of safety for therapeutic applications.^14^ However, application in HSCs presents distinct challenges, as engineered epigenetic states must be stably maintained through the extensive cell division during both self-renewal and hematopoietic differentiation. While a prior study reported that targeted partial promoter methylation at the *CDKN2B* (p15) locus in human hematopoietic stem and progenitor cells (HSPCs) persisted during myeloid differentiation *in vitro* and in xenotransplanted mice,^20^ the comprehensive characterization of targeted epigenome silencing durability during HSC self-renewal and multilineage differentiation, its reversibility, and therapeutic potential in human hematopoiesis remains largely unexplored.

Thrombosis, the formation of a clot within blood vessels, is a major cause of life-threatening conditions such as ischemic heart disease and strokes that are estimated to impact over 40 million people around the world each year.^21,22^ A mainstay to prevent and treat these clots involves blockade of platelet activation through the use of antiplatelet therapies, such as aspirin and clopidogrel.^23–26^ Unfortunately, these therapies require lifelong daily administration, which is often complicated by poor compliance and some patients can develop resistance to these therapies, limiting the effectiveness of clot prevention.^27,28^ Moreover, platelet function studies have shown that even among patients receiving standard antiplatelet therapy, a subset have persistently high on-treatment platelet reactivity, which is associated with an increased risk of recurrent thrombotic events.^29,30^ Therefore, there is a need to develop improved antiplatelet therapies that are durable and broadly effective. Platelets are derived from megakaryocytes that differentiate from HSCs and downstream progenitors.^31^ Thus, platelet biology provides a relevant proof-of-principle application through which to assess the durability and functional consequences of epigenetic programming in HSCs and their differentiated progeny.

Here, we optimize and apply highly efficient epigenetic silencing in human HSCs. As an initial proof-of-principle target, we chose the high-copy platelet fibrinogen receptor integrin β3 (*ITGB3*), which is required for platelet aggregation and is a key target of antithrombotic therapies.^32–34^ We demonstrate robust *ITGB3* silencing by CHARM that persists through HSC self-renewal and megakaryocytic differentiation, resulting in significantly reduced αIIbβ3-dependent aggregation of *in vitro*-derived platelets. Silencing remained stable in engrafted human HSPCs four months after xenotransplantation into NBSGW mice. Targeted demethylation partially reversed *ITGB3* silencing, and targeting additional platelet-associated genes demonstrated the broader applicability of this approach. These results establish durable and reversible epigenetic programming of HSCs as a platform for modulation of platelet function, highlighting its potential for thrombosis prevention, while also motivating future investigation of its applicability to other diseases rooted in the HSC compartment.

## Results

### Optimization of approaches for programmable epigenetic silencing in human HSCs

We first sought to optimize conditions for applying targeted programmable epigenetic silencing in human HSCs and their progeny using RNA delivery (**Figure 1A**). CD34^+^ HSPCs that were mobilized from healthy adult donors were electroporated with mRNAs encoding CRISPR interference (CRISPRi), CRISPRoff, or CHARM (**Figure S1A**), together with a single guide RNA (sgRNA) targeting the promoter of *CD47*, selected for its annotated promoter CpG island, cell surface localization, broad hematopoietic expression, and functional neutrality in our *in vitro* culture conditions. Cells were maintained in HSPC expansion culture, and CD47 expression was monitored by flow cytometry over time. CRISPRi initially abolished CD47 expression in ~90% of cells, but the effect progressively diminished over several days in culture (**Figures 1B-C**), consistent with the short-lived nature of RNA delivery and the generally non-heritable characteristics of histone modifications. In contrast, DNA methylation-based silencers, CRISPRoff and CHARM, induced efficient (~80-95% CD47^−^ cells) and persistent silencing in HSPCs throughout an extended culture (**Figures 1B-C**). Similar repression was observed when the analysis was restricted to the CD34^+^CD45RA^−^CD90^+^ HSC-enriched subset, indicating effective gene silencing in populations enriched for *bona fide* HSCs (**Figures 1C** and **S1B-C**). We selected CHARM for subsequent experiments due to its smaller size and favorable safety profile, which align with therapeutic development.^15^ Consistent with an epigenetic mechanism of silencing, CHARM-mediated repression was accompanied by increased DNA methylation surrounding the *CD47* transcription start site (TSS) and concomitantly reduced mRNA expression (**Figures 1D** and **S1D**).

**Figure 1.**
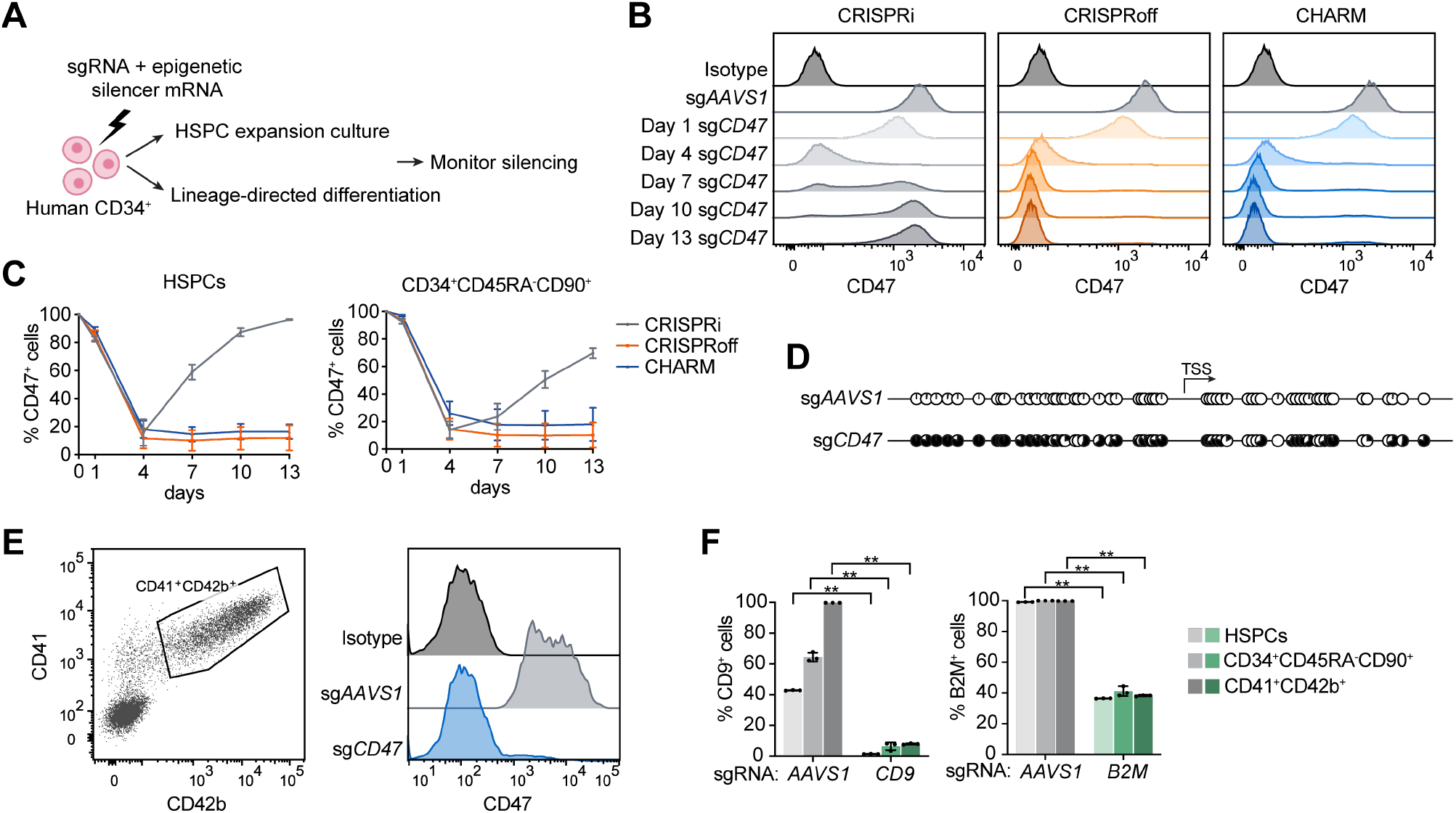
CHARM enables durable epigenetic silencing in human HSPCs. **(A)** Schematic overview of the workflow used to establish epigenetic silencing in primary human CD34^+^ HSPCs. **(B)** Representative flow cytometry histograms of surface CD47 expression over time in HSPCs edited with CRISPRi, CRISPRoff, or CHARM targeting the *CD47* promoter. Isotype staining and *AAVS1*-targeted cells serve as controls. **(C)** Quantification of the percentage of CD47^+^ cells over time in bulk HSPCs (left) and within phenotypically defined CD34^+^CD45RA^−^CD90^+^ HSCs (right) following CRISPRi, CRISPRoff, or CHARM targeting of the *CD47* promoter (n = 2 donors). **(D)** Targeted bisulfite sequencing displaying CpG methylation patterns near the *CD47* TSS in HSPCs 13 days after editing with CHARM targeting the *CD47* promoter or AAVS1 as a control. Each circle represents an individual CpG site; filled circles indicate methylated CpGs. *P* < 0.0001. **(E)** Left, representative flow cytometry gating strategy used to define CD41^+^CD42b^+^ mature megakaryocytes derived from edited HSPCs. Right, representative flow cytometry histograms of surface CD47 expression in these cells following CHARM targeting of *CD47* promoter or *AAVS1*. **(F)** Quantification of the percentage of CD9^+^ (left) and B2M^+^ (right) cells across bulk HSPCs and CD34^+^CD45RA^−^CD90^+^-defined HSCs at 13 days after editing, and CD41^+^CD42b^+^ megakaryocytes after 12 days of differentiation, following CHARM targeting in HSPCs (n = 3). All data are presented as mean ± SD, significance is indicated as \**P* < 0.05, \*\**P* < 0.01, or n.s. not significant.

To assess whether epigenetic silencing remained stable after differentiation of HSCs, CHARM-edited HSPCs were differentiated toward the megakaryocytic and erythroid lineages. We found that *CD47* repression was stably maintained at the endpoints of both trajectories, with ~95% silencing in CD41^+^CD42b^+^ mature megakaryocytes (MKs) and ~99% silencing in erythroid cultures composed predominantly of intermediate (CD71^+^CD235^+^) and late (CD71^−^CD235^+^) erythroblasts (**Figures 1E** and **S1E-F**), indicating preserved repression through lineage commitment and terminal maturation. Importantly, despite known alterations in global DNA methylation during hematopoietic differentiation^35–38^, these results support the stability and effectiveness of this approach.

We then evaluated the generality of CHARM-mediated silencing by targeting two additional surface genes with distinct expression profiles, *CD9* and *B2M*. Both loci exhibited robust and durable repression across HSPCs, HSC-enriched populations, and differentiated MKs, albeit with varying silencing efficiencies (>90% CD9^−^ cells and ~60% B2M^−^ cells) (**Figure 1F**). Together, these results establish CHARM as an efficient and durable epigenome silencing platform for human HSCs that remains stable upon differentiation.

### Effective and specific *ITGB3* silencing by CHARM

Persistent gene silencing in human HSCs and their differentiated progeny could enable sustained modulation of hematopoietic cell function. We reasoned that this property could be leveraged to silence genes essential for platelet adhesion, activation, and aggregation, with the potential to reduce thrombosis risk. A key target is the integrin αIIbβ3 (encoded by *ITGA2B* and *ITGB3*), a central mediator of platelet aggregation that acts through fibrinogen binding.^32,33^ Loss-of-function variants in *ITGA2B* and *ITGB3* cause Glanzmann thrombasthenia, a bleeding disorder that lacks other phenotypic manifestations; clinically, αIIbβ3 inhibitors are used to prevent thrombotic complications, particularly during acute interventions.^39,40^ Extensive prior work has defined the central role of *ITGB3* in platelet aggregation, and its CpG-rich promoter architecture makes it particularly well suited for CHARM-mediated methylation, which collectively nominate *ITGB3* as an ideal proof-of-principle target.^32–34,41^ We hypothesized that CHARM-induced methylation at the *ITGB3* promoter would persist through HSC self-renewal and megakaryocytic differentiation, leading to stable silencing in MKs and derived platelets and, consequently, impaired aggregation (**Figure 2A**).

**Figure 2.**
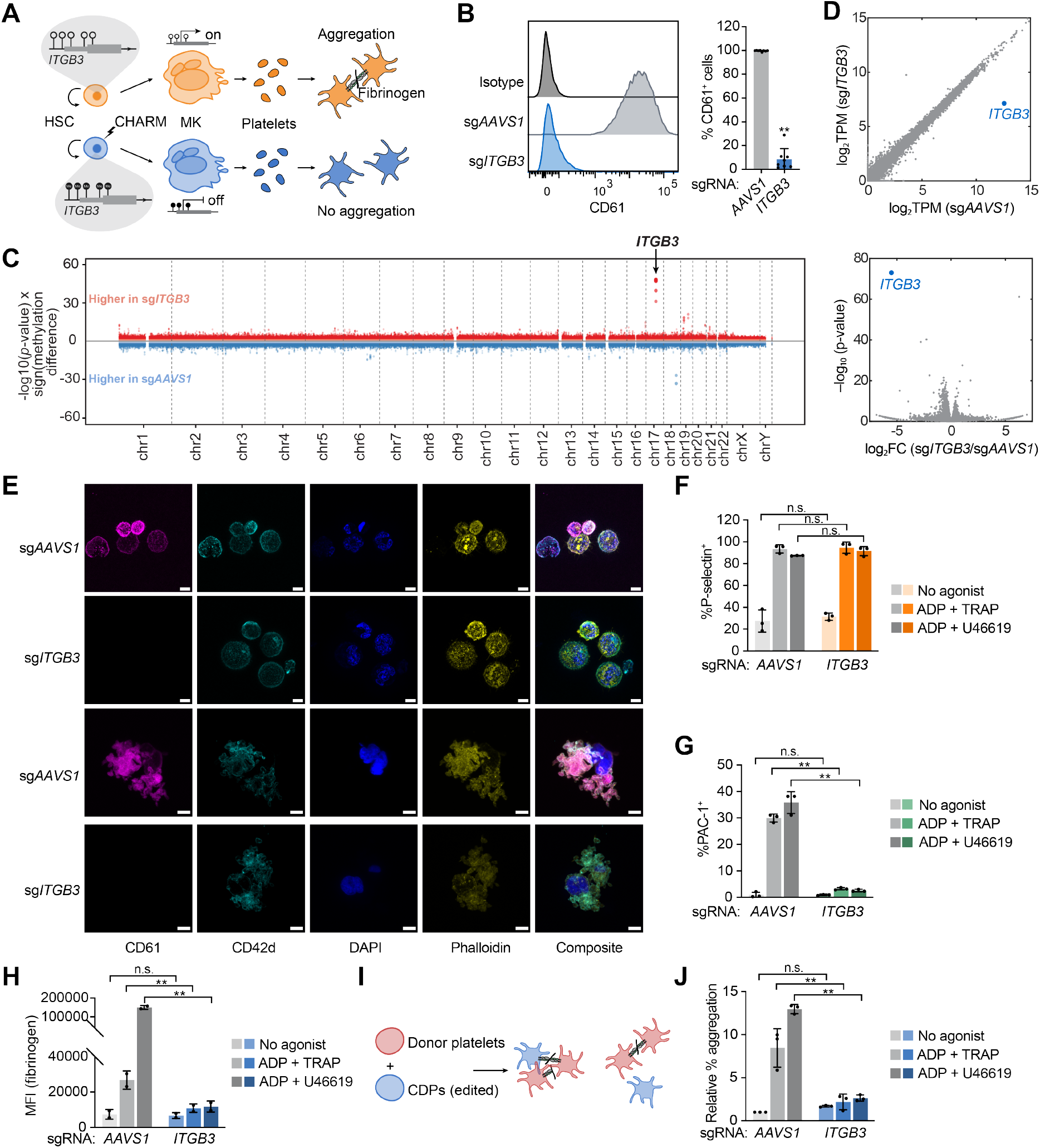
Robust and specific silencing of *ITGB3* by CHARM persists through megakaryocytic differentiation and impairs αIIbβ3-dependent platelet function. **(A)** Schematic illustrating the strategy for CHARM-mediated silencing of *ITGB3* in HSPCs. Targeted methylation established in HSPCs is maintained through self-renewal, megakaryocytic differentiation, and platelet formation, resulting in reduced αIIbβ3 expression and platelet aggregation. **(B)** Left, representative flow cytometry histograms of CD61 (*ITGB3*) surface expression in mature megakaryocytes differentiated from HSPCs following CHARM targeting of the *ITGB3* promoter or *AAVS1*. Right, quantification of the percentage of CD61^+^ cells among mature megakaryocytes (n = 8 donors). **(C)** Manhattan plot displaying differentially methylated regions between HSPCs edited with CHARM targeting the *ITGB3* promoter or *AAVS1* as a control, analyzed by WGBS. Red dots indicate CpGs with increased DNA methylation in *ITGB3*-targeted cells relative to *AAVS1* controls, whereas blue dots indicate CpGs with decreased methylation. The arrow denotes the genomic position of *ITGB3*. **(D)** RNA sequencing plots of differentiated megakaryocytes edited with CHARM targeting the *ITGB3* promoter or *AAVS1* as a control. Top, scatter plot showing transcript abundance (log2 TPM) for each gene in the two experimental groups, with *ITGB3* highlighted in blue. Bottom, volcano plot showing differential gene expression between conditions, with *ITGB3* highlighted in blue. **(E)** Representative immunofluorescence images of megakaryocytes and proplatelets derived from HSPCs edited with CHARM targeting the *ITGB3* promoter or *AAVS1* as a control. Cells were stained for CD61, CD42d, DAPI, and F-actin (phalloidin). Scale bars, 10 µm. **(F)** Quantification of P-selectin induction in CDPs generated from HSPCs edited with CHARM targeting the *ITGB3* promoter or *AAVS1* as a control under no-agonist conditions or following stimulation with ADP + TRAP or ADP + U46619 (n = 3 donors). **(G)** Quantification of PAC-1 binding in CDPs generated from HSPCs edited with CHARM targeting the *ITGB3* promoter or *AAVS1* as a control under no-agonist conditions or following stimulation with ADP + TRAP or ADP + U46619 (n = 3 donors). **(H)** Quantification of fibrinogen binding, measured as fibrinogen MFI, in CDPs generated from HSPCs edited with CHARM targeting the *ITGB3* promoter or *AAVS1* as a control under no-agonist conditions or following stimulation with ADP + TRAP or ADP + U46619 (n = 2 donors). **(I)** Schematic of the platelet aggregation assay assessing the ability of CDPs generated from edited HSPCs to aggregate with donor platelets. **(J)** Quantification of relative platelet aggregation in CDPs generated from HSPCs edited with CHARM targeting the *ITGB3* promoter or *AAVS1* as a control, measured as the percentage of double-positive platelet-CDP events normalized to total CDP events under no-agonist conditions or following stimulation with ADP + TRAP or ADP + U46619 (n = 3 donors, platelets from 2 donors). All data are presented as mean ± SD, significance is indicated as \**P* < 0.05, \*\**P* < 0.01, or n.s. not significant.

In this context, we first examined whether *ITGB3* could be silenced by CHARM in HSPCs and maintained through megakaryocytic differentiation. CD34^+^ HSPCs were electroporated with CHARM mRNA together with an sgRNA targeting either the *AAVS1* control locus or the *ITGB3* promoter. Following differentiation, the latter induced robust silencing of *ITGB3*, with an average of ~90% CD61 (encoded by *ITGB3*) negative cells among mature MKs (**Figures 2B** and **S2A)**. In line with promoter-targeted epigenetic repression, we observed a significant increase in DNA methylation near the *ITGB3* TSS and a reduction in *ITGB3* mRNA expression (**Figures S2B-C**).

To assess the specificity of CHARM-mediated DNA methylation, we performed whole-genome bisulfite sequencing (WGBS) on modified HSPCs. Compared with *AAVS1* targeting controls, the most prominent increase in DNA methylation was observed at the targeted *ITGB3* locus on chromosome 17 (**Figure 2C**). Consistent with prior reports, in addition to focal methylation at the sgRNA targeting site, DNA methylation spread across the CpG-rich region surrounding the *ITGB3* promoter (~1.5 kb) (**Figure S2D**). WGBS also identified a differentially methylated region (DMR) on chromosome 18 with higher methylation in the *AAVS1* control. However, targeted bisulfite sequencing of this region showed comparable methylation in unedited, CHARM-*AAVS1*, and CHARM-*ITGB3* HSPCs, indicating that the difference was not reproducible and likely reflected low WGBS coverage at this locus (**Figure S2E**). Off-target DMRs identified in HSPCs were subsequently hierarchically annotated based on genomic context (**Figure S2F**). Notably, *in silico*-predicted sgRNA off-target sites did not overlap these regions. Although seven off-target DMRs were annotated as promoter-associated, inspection of these loci revealed that differential methylation tended to involve limited regions and rarely overlapped core promoter CpG islands or TSSs (**Figure S2G**). Notably, genes associated with these promoter-proximal off-target DMRs did not exhibit significant transcriptional changes in edited HSPCs, nor was there evidence of widespread transcriptional perturbation (**Figures S2H-I**). To further examine transcriptional consequences of CHARM-mediated *ITGB3* silencing following megakaryocytic differentiation, RNA sequencing was performed on differentiated and sorted CD42b^+^ MKs. We found that gene repression was highly specific to *ITGB3*, while other transcripts, including those associated with off-target DMRs identified by WGBS, remained largely unchanged. The only notable exception was up-regulation of the neighboring gene of *ITGB3, MYL4*, likely reflecting local chromatin remodeling at the silenced locus, as has been observed with DNA methylation changes occurring during hematopoietic differentiation (**Figure 2D** and **S2J**).^38^

### Disruption of platelet aggregation by CHARM-mediated *ITGB3* silencing

To further assess *ITGB3* silencing during MK maturation and proplatelet formation, we visualized the cells through immunofluorescence imaging. We observed mature, round, CD42-positive, polyploid MKs and readily detected proplatelet-forming structures, confirming the presence of morphologically conventional MKs. In parallel, flow cytometric analysis of DNA content revealed comparable ploidy distributions between control and CHARM *ITGB3*-edited MKs (**Figure S3A**). While control cells exhibited clear CD61 labeling, CHARM *ITGB3*-edited cells showed near-complete loss of CD61 signal, indicating efficient suppression of *ITGB3* expression (**Figures 2E** and **S3B**).

We next sought to examine the functional consequences of CHARM-mediated *ITGB3* silencing in derived platelets (**Figure S3C**). Although culture-derived platelets (CDPs) produced *in vitro* from differentiated MKs tend to be partially pre-activated and have reduced function compared to platelets produced *in vivo*, they can be valuable surrogates to examine physiological responses.^42–45^ Following stimulation with ADP together with the PAR1 agonist peptide TRAP or the thromboxane A_2_ analog U46619, both control and *ITGB3*-silenced CDPs exhibited similar robust upregulation of surface P-selectin, indicating that agonist-induced platelet activation remain largely intact (**Figures 2F** and **S3D**). In contrast, PAC-1 binding was markedly reduced in *ITGB3*-silenced CDPs, consistent with the loss of active αIIbβ3 integrin on the platelet surface (**Figures 2G** and **S3E**). We additionally assessed agonist-induced fibrinogen binding and observed a marked reduction in *ITGB3*-silenced CDPs compared with controls (**Figures 2H and S3F**). Together, these findings are consistent with the specific targeting of *ITGB3*, resulting in selective impairment of αIIbβ3-dependent platelet responses, while preserving other aspects of agonist-induced platelet activation.

To further determine whether these translated into impaired platelet aggregation, we employed a flow cytometry-based assay to assess the ability of CDPs generated from edited cells^45^ to aggregate with healthy donor platelets (**Figure 2I**). Donor platelets and CDPs were differentially labeled, mixed, and stimulated with agonists before fixation and flow cytometry (**Figure S3G**). Double-positive events, representing platelet-CDP aggregates, were quantified as a measure of CDP aggregation capacity. While control CDPs exhibit robust aggregation upon stimulation with either agonist combination, those derived from *ITGB3*-silenced cells displayed a pronounced defect in forming aggregates with donor platelets, consistent with the essential role of αIIbβ3 in platelet-platelet interactions that are induced by a variety of platelet activators (**Figures 2J** and **S3H**). Collectively, these results demonstrate that silencing of *ITGB3* by CHARM persists through MK differentiation, generating platelets that lack this key fibrinogen receptor, thereby impairing platelet aggregation.

### Persistence of *ITGB3* silencing by CHARM through HSC self-renewal

Beyond differentiation, effective HSC-based approaches require that editing outcomes be maintained as HSCs undergo self-renewal throughout the lifespan. We therefore sought to rigorously examine persistence of silencing during HSC self-renewal and long-term culture. To track cell division, CHARM-edited HSPCs were labeled with carboxyfluorescein succinimidyl ester (CFSE). Division of phenotypically defined CD34^+^CD45RA^−^CD90^+^ HSC-enriched populations was observed over five days of expansion culture (**Figures 3A** and **S4A**). We found that DNA methylation surrounding the *ITGB3* TSS was well preserved in these cells, indicating maintenance of targeted DNA methylation through division of primitive HSPCs (**Figure 3B**). These sorted HSCs were further subjected to megakaryocytic differentiation, during which *ITGB3* silencing was robustly retained in mature MKs, as assessed by a lack of CD61 expression, demonstrating transmission of silencing from dividing HSCs to their differentiated progeny (**Figure 3C**). Apoptosis, cell viability, and the maintenance of phenotypically defined short-and long-term repopulating (ST/LT) HSCs were not affected, consistent with the specificity of editing (**Figures S4B-C**).

**Figure 3.**
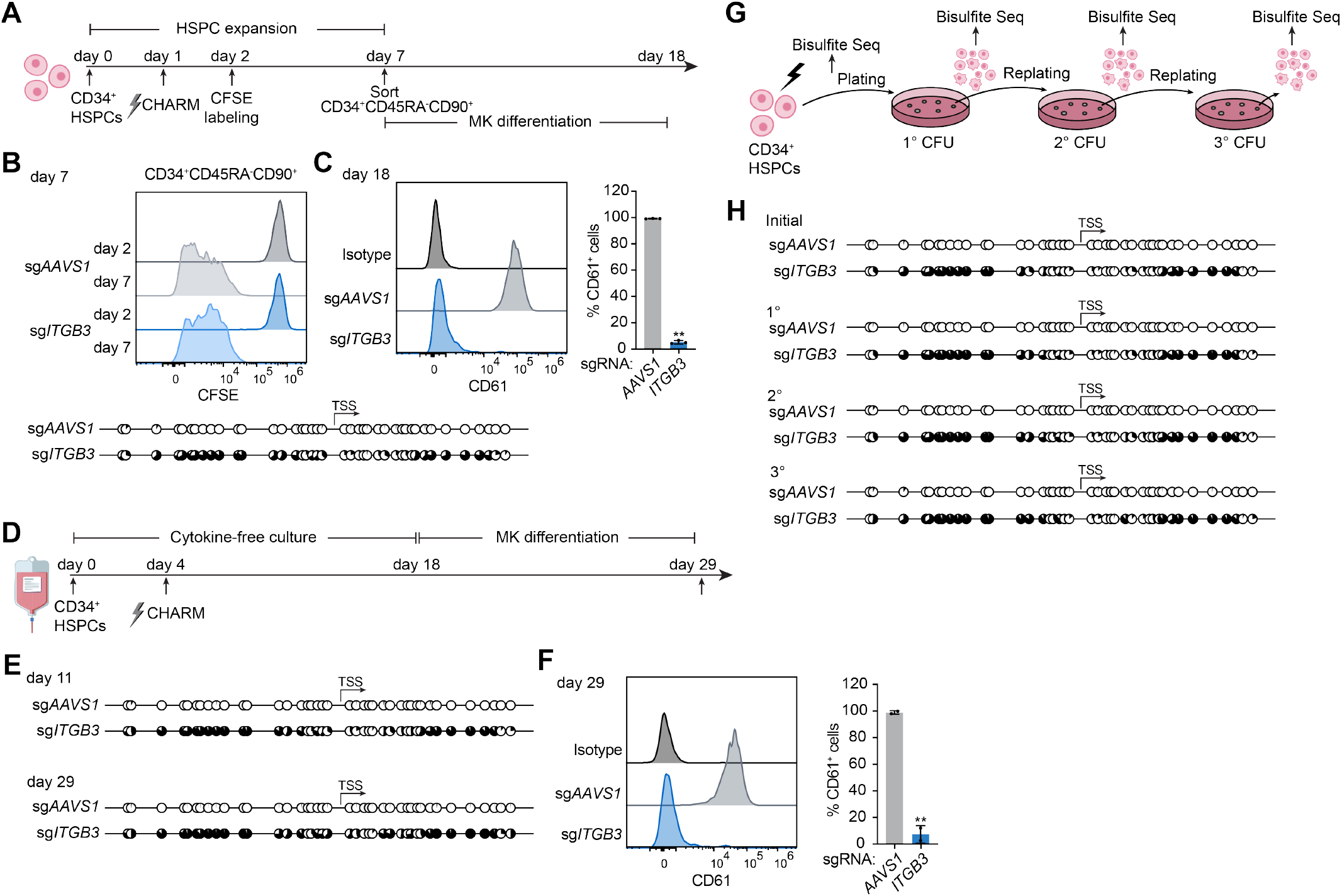
CHARM-mediated *ITGB3* silencing persists through HSC self-renewal. **(A)** Schematic overview of the experimental design for CHARM editing, tracking and sorting of CD34^+^CD45RA^−^CD90^+^ cells, and subsequent megakaryocytic differentiation. **(B)** Top, representative CFSE dilution profiles of CD34^+^CD45RA^−^CD90^+^ cells at days 2 and 7 following CHARM targeting of the *ITGB3* promoter or *AAVS1* as a control. Bottom, targeted bisulfite sequencing displaying CpG methylation patterns near the *ITGB3* TSS in the sorted CD34^+^CD45RA^−^CD90^+^ cells. Each circle represents an individual CpG site; filled circles indicate methylated CpGs. **(C)** Left, representative flow cytometry histograms of CD61 (*ITGB3*) surface expression in megakaryocytes differentiated from sorted CD34^+^CD45RA^−^CD90^+^ cells following CHARM targeting of the *ITGB3* promoter or *AAVS1* as a control. Right, quantification of the percentage of CD61^+^ cells among differentiated megakaryocytes (n = 3 donors). **(D)** Schematic overview of the experimental design for extended cytokine-free culture and subsequent megakaryocytic differentiation following CHARM editing of HSPCs. **(E)** Targeted bisulfite sequencing displaying CpG methylation patterns near the *ITGB3* TSS in HSPCs at day 11 and day 29 following CHARM targeting of the *ITGB3* promoter or AAVS1 as a control. Each circle represents an individual CpG site; filled circles indicate methylated CpGs. **(F)** Left, representative flow cytometry histograms of CD61 (*ITGB3*) surface expression in megakaryocytes differentiated from HSPCs after extended cytokine-free culture following CHARM targeting of the *ITGB3* promoter or *AAVS1* as a control. Right, quantification of the percentage of CD61^+^ cells among differentiated megakaryocytes (n = 2 donors). **(G)** Schematic overview of the experimental design for serial CFU replating and targeted bisulfite sequencing following CHARM editing of HSPCs. **(H)** Targeted bisulfite sequencing displaying CpG methylation patterns near the *ITGB3* TSS in HSPCs after CHARM editing and in cells derived from primary (1°), secondary (2°), and tertiary (3°) CFU replatings following targeting of the *ITGB3* promoter or AAVS1 as a control. Each circle represents an individual CpG site; filled circles indicate methylated CpGs. All data are presented as mean ± SD, significance is indicated as \**P* < 0.05, \*\**P* < 0.01, or n.s. not significant.

To enable longer-term evaluation of silencing stability, edited cells were maintained in a chemically defined, cytokine-free culture which maximizes HSC self-renewal *in vitro* for three weeks prior to megakaryocytic differentiation (**Figure 3D**).^46,47^ Throughout this extended culture, we found that DNA methylation at the *ITGB3* TSS remained stable (**Figure 3E** and **S4D**). At the endpoint of differentiation, loss of CD61 expression was still observed in mature MKs, indicating that *ITGB3* silencing was maintained after prolonged culture (**Figure 3F**).

Lastly, to assess maintenance of DNA methylation during functional HSPC self-renewal and multilineage differentiation, we performed serial replating assays. Edited HSPCs were plated in methylcellulose, and colonies were harvested and replated sequentially to generate secondary and tertiary colonies that depend upon progenitor and stem cell self-renewal (**Figures 3G** and **S4E**). Targeted DNA methylation near the *ITGB3* TSS was examined after each replating, and was faithfully maintained through tertiary replating (**Figure 3H** and **S4F**). These results indicate stable transmission of CHARM-induced DNA methylation across repeated cycles of HSPC self-renewal and differentiation.

### Sustained silencing during long-term engraftment in mice

To further assess the long-term persistence and functional output of edited human HSCs *in vivo*, we performed xenotransplantation of edited human CD34^+^ HSPCs into *Kit* mutant and immunodeficient NOD.Cg-*Kit*^*W-41J*^*Tyr*^+^*Prkdc*^*scid*^*Il2rg*^*tm1Wjl*^/ThomJ (NBSGW) mice (**Figure 4A**). 4 months post transplantation, after long-term engraftment by human HSCs is well established, bone marrow was harvested for analysis.

**Figure 4.**
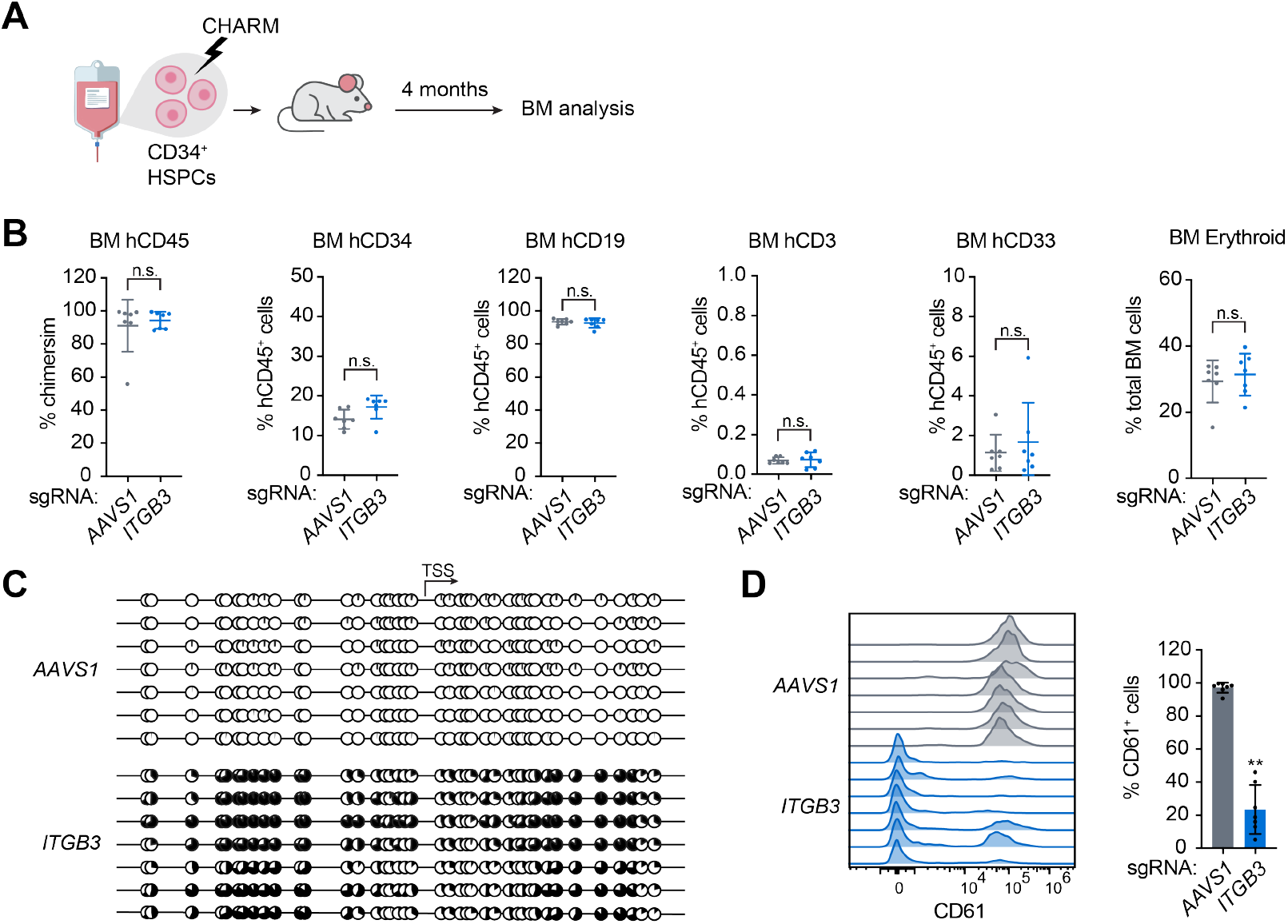
*ITGB3* silencing by CHARM is maintained following long-term engraftment in xenotransplanted mice. **(A)** Schematic overview of xenotransplantation of CHARM-edited human CD34^+^ HSPCs into NBSGW mice and subsequent analysis 16 weeks after transplantation. (**B)** Quantification of human hematopoietic engraftment in the bone marrow of recipient mice 16 weeks after transplantation, assessed by overall chimerism (% hCD45^+^ cells among total human plus mouse CD45^+^ cells), and frequency of human CD34^+^ compartment, B (hCD19^+^), T (hCD3^+^), myeloid (hCD33^+^) (expressed as a percentage of human hCD45^+^ cells), and erythroid populations (CD71^+^ and/or CD235a^+^ cells expressed as a percentage of total bone marrow cells). **(C)** Targeted bisulfite sequencing displaying CpG methylation patterns near the *ITGB3* TSS in human CD34^+^ cells isolated from transplanted mice 16 weeks after engraftment. Each circle represents an individual CpG site; filled circles indicate CpGs with high DNA methylation. **(D)** Representative flow cytometry histograms of CD61 (*ITGB3*) surface expression in megakaryocytes differentiated from human CD34^+^ cells recovered from mice transplanted with CHARM-edited HSPCs targeting the *ITGB3* promoter or *AAVS1* as a control. All data are presented as mean ± SD, significance is indicated as \**P* < 0.05, \*\**P* < 0.01, or n.s. not significant.

Robust and comparable human hematopoietic engraftment was observed in recipient mice transplanted with CHARM-edited HSPCs targeting either the *AAVS1* locus or the *ITGB3* promoter, as assessed by the frequency of hCD45^+^ cells in the bone marrow (**Figure 4B**). Similarly, the contribution of the CD34^+^ compartment and major hematopoietic lineages, including B (CD19^+^), T (CD3^+^), myeloid (CD33^+^), and erythroid populations, was comparable between the two groups (**Figures 4B** and **S5A-C**), demonstrating a lack of undesired effects on HSPC engraftment or lineage output.

To examine whether CHARM-mediated epigenetic modification at the *ITGB3* locus was maintained in engrafting human progenitors after self-renewal of HSCs post-transplant and residence in the bone marrow niche, we performed bisulfite sequencing on human CD34^+^ cells isolated from the bone marrows of transplanted mice. Sustained DNA methylation near the *ITGB3* TSS was observed in all mice in the *ITGB3* targeting group (**Figure 4C** and **S5D**). Given that few human platelets or megakaryocytes were generated in these mice, we subjected CD34^+^ cells from long-term engrafted mice to megakaryocytic differentiation, and found that *ITGB3* silencing was robustly retained in mature MKs, with on average ~80% of cells lacking CD61 expression (**Figure 4D**).

### Reversibility of silencing through targeted demethylation

One key feature of epigenome editing is its reversibility, enabling restoration of gene expression if required in a therapeutic context.^14,17^ We therefore tested whether targeted demethylation using TET1-dCas9 could reverse CHARM-mediated repression by erasing methylation marks at the silenced promoter (**Figure 5A**). To this end, CHARM-edited HSPCs were maintained in cytokine-free culture for eight days and then a second electroporation was performed with mRNA encoding TET1-dCas9, together with the sgRNAs targeting either the *AAVS1* control locus or the *ITGB3* promoter (**Figure 5B**). Following megakaryocytic differentiation, the latter induced significant demethylation near the *ITGB3* TSS (**Figure 5C**). Correspondingly, surface CD61 expression was restored (**Figure 5D**). These results demonstrate that targeted demethylation enables controlled reversal of silencing in human HSPCs.

**Figure 5.**
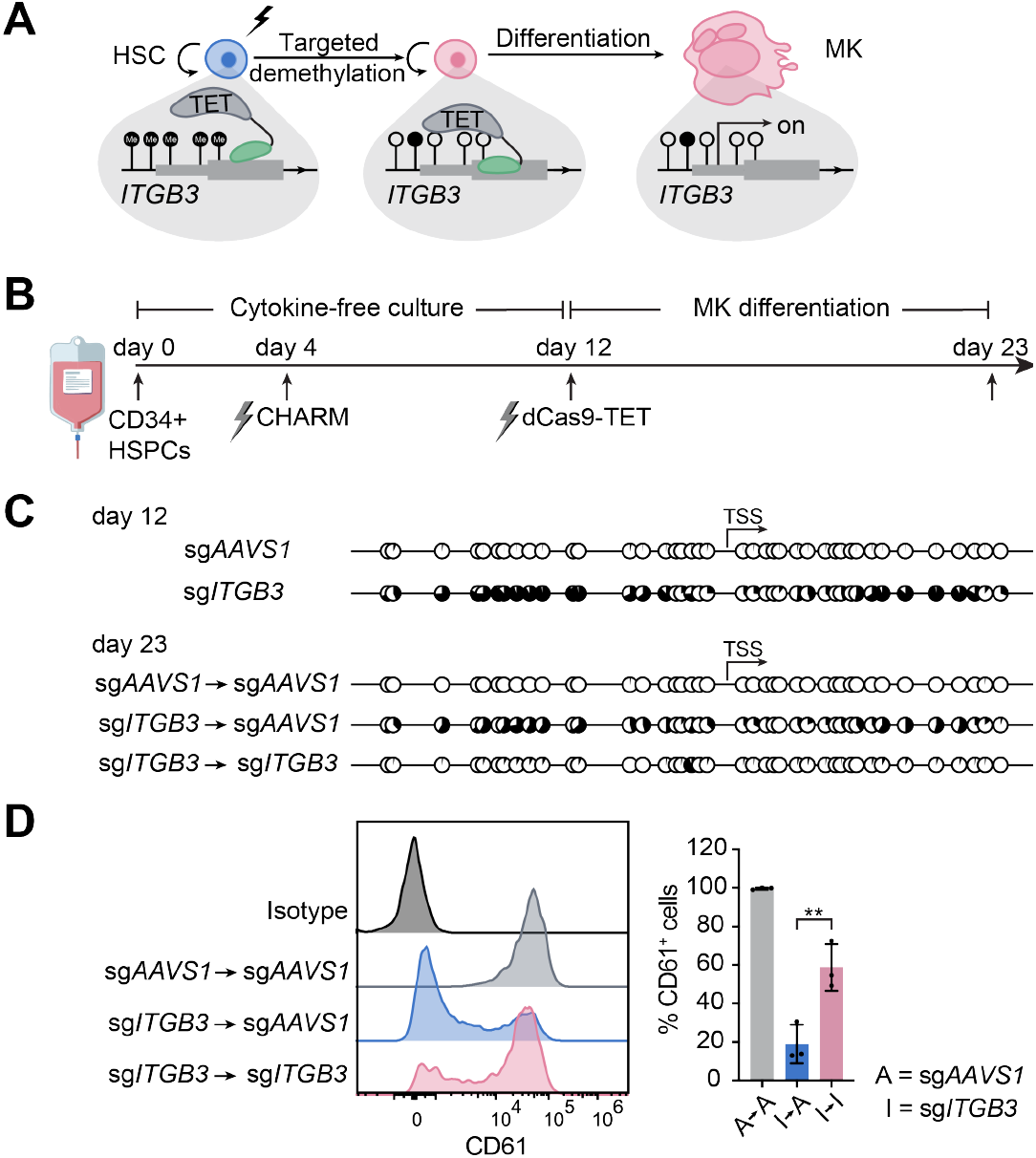
Targeted demethylation reverses CHARM-mediated *ITGB3* silencing. **(A)** Schematic illustrating targeted demethylation of the *ITGB3* promoter in HSCs using dCas9-TET following CHARM-mediated silencing, and subsequent megakaryocytic differentiation. **(B)** Schematic overview of the experimental design for sequential CHARM-mediated methylation and dCas9-TET-mediated demethylation in HSPCs, followed by megakaryocytic differentiation. **(C)** Representative targeted bisulfite sequencing displaying CpG methylation patterns near the *ITGB3* TSS in HSPCs at day 12 (before targeted demethylation) and day 23 (after targeted demethylation and megakaryocytic differentiation), following CHARM and dCas9-TET targeting as indicated. Each circle represents an individual CpG site; filled circles indicate methylated CpGs. (*P* < 0.0001 for sgITGB3→sgAAVS1 versus sgITGB3→sgITGB3 at day 23).. **(D)** Left, representative flow cytometry histograms of CD61 (*ITGB3*) surface expression in megakaryocytes differentiated from HSPCs subjected to sequential CHARM and dCas9-TET targeting as indicated. Right, quantification of the percentage of CD61^+^ cells among megakaryocytes (n = 3 donors). All data are presented as mean ± SD, significance is indicated as \**P* < 0.05, \*\**P* < 0.01, or n.s. not significant.

### Broad applicability of CHARM to modulate platelet function

While the integrin αIIbβ3 represents a valuable initial proof-of-principle target to demonstrate the effectiveness of CHARM to prevent platelet aggregation after HSC editing, additional platelet targets act at distinct stages of thrombus formation and may carry different bleeding risks and safety profiles. We therefore asked whether CHARM-mediated silencing in HSPCs could be extended to additional genetically supported platelet-associated genes as candidate targets for modulating platelet function (**Figure 6A**).

**Figure 6.**
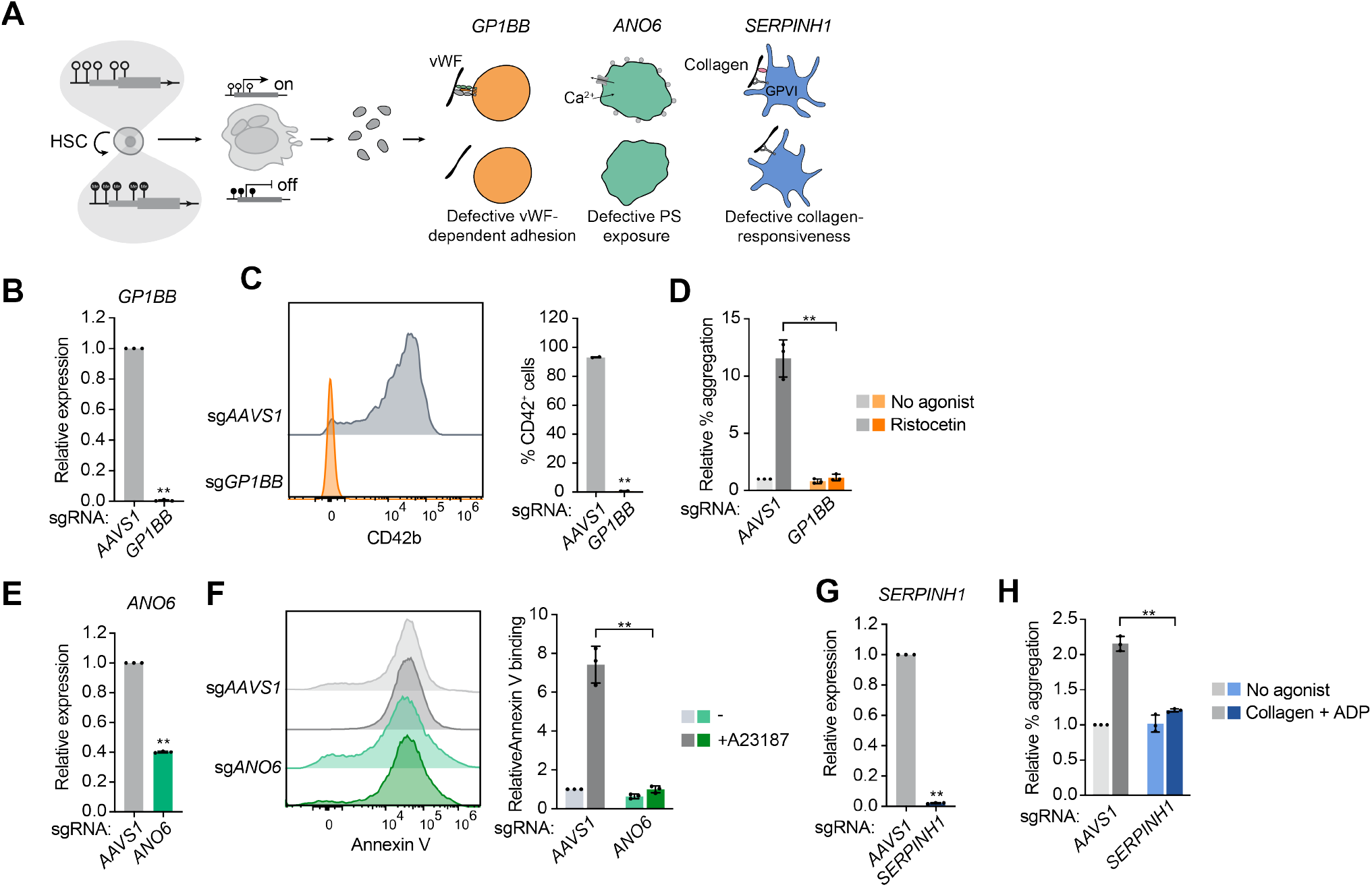
CHARM is broadly applicable for epigenetic silencing of multiple platelet genes in HSPCs. (**A)** Schematic illustrating CHARM-mediated epigenetic silencing in HSPCs and its application to distinct platelet genes acting at different stages of thrombus formation, including *GP1BB, ANO6*, and *SERPINH1*. **(B)** Quantification of *GP1BB* mRNA expression measured by RT-qPCR in differentiated megakaryocytes following CHARM targeting of the *GP1BB* promoter or *AAVS1* as a control (n = 3 donors). **(C)** Left, representative flow cytometry histograms of CD42b surface expression in CD41^+^GP6^+^ megakaryocytes differentiated from HSPCs following CHARM targeting of the *GP1BB* promoter or *AAVS1*. Right, quantification of the percentage of CD42b^+^ cells among megakaryocytes (n = 2 donors). **(D)** Quantification of relative platelet aggregation in CDPs generated from HSPCs edited with CHARM targeting the *GP1BB* promoter or *AAVS1* as a control, measured as the percentage of double-positive platelet-CDP events normalized to total CDP events under no-agonist conditions or following stimulation with ristocetin ristocetin (n = 3 donors, platelets from 2 donors). **(E)** Quantification of *ANO6* mRNA expression measured by RT-qPCR in differentiated CD41^+^CD42b^+^ megakaryocytes following CHARM targeting of the *ANO6* promoter or *AAVS1* as a control (n = 3 donors). **(F)** Left, representative flow cytometry histograms of Annexin V binding in CDPs generated from HSPCs edited with CHARM targeting the *ANO6* promoter or *AAVS1* under baseline conditions or following stimulation with the Ca2^+^ ionophore A23187. Right, quantification of relative Annexin V binding (n = 3 donors). **(G)** Quantification of *SERPINH1* mRNA expression measured by RT-qPCR in differentiated CD41^+^CD42b^+^ megakaryocytes following CHARM targeting of the *SERPINH1* promoter or *AAVS1* as a control (n = 3 donors). **(H)** Quantification of relative platelet aggregation in CDPs generated from HSPCs edited with CHARM targeting the *SERPINH1* promoter or *AAVS1* as a control, measured as the percentage of double-positive platelet-CDP events normalized to total CDP events under no-agonist conditions or following stimulation with collagen + ADP (n = 3 donors, platelets from 2 donors). All data are presented as mean ± SD, significance is indicated as \**P* < 0.05, \*\**P* < 0.01, or n.s. not significant.

We first targeted *GP1BB*, which encodes a component of the GPIb-IX-V complex that mediates von Willebrand factor (vWF)-dependent platelet adhesion under high shear conditions.^48^ This target has been validated through loss-of-function human genetic mutations in the gene that result in the bleeding disorder Bernard-Soulier syndrome.^49^ CHARM targeting of the *GP1BB* promoter in HSPCs resulted in increased DNA methylation at the targeted locus in differentiated MKs (**Figure S6A**), accompanied by near-complete repression of *GP1BB* mRNA expression (**Figure 6B**). This silencing led to loss of surface CD42b (GPIbα), consistent with destabilization of the GPIb complex (**Figure 6C**). Functionally, CDPs generated from *GP1BB*-silenced HSPCs exhibited a profound defect in ristocetin-induced aggregation with donor platelets, demonstrating impaired GPIb-vWF-dependent platelet aggregation (**Figures 6D** and **S6B**).

We next tested CHARM-mediated silencing of *ANO6*, which encodes a Ca^2+^-activated phospholipid scramblase that is essential for platelet procoagulant activity through phosphatidylserine (PS) exposure.^50–52^ Loss-of-function mutations in this gene cause Scott syndrome, a mild bleeding disorder.^50^ Editing of *ANO6* by CHARM led to a partial but substantial reduction (~60%) in *ANO6* mRNA expression in differentiated MKs, with corresponding increased DNA methylation near the TSS (**Figures 6E** and **S6C**). At baseline, CDPs displayed some Annexin V binding (indicating surface PS exposure), consistent with partial membrane remodeling and pre-activation commonly observed in platelets derived from culture. However, stimulation with the Ca^2+^ ionophore A23187 induced a pronounced loss of the PS-negative population in control CDPs, whereas this response was markedly attenuated in *ANO6*-silenced CDPs (**Figure 6F**), indicating impaired Ca^2+^-triggered PS externalization.

Finally, we examined the potential of silencing *SERPINH1* (HSP47), a chaperone implicated in platelet activation downstream of collagen engagement.^53,54^ Interestingly, reduced *SERPINH1* expression in platelets has been suggested to prevent thrombosis in hibernating animals, such as bears, and in immobile humans.^53^ *SERPINH1* targeting by CHARM in HSPCs led to robust DNA methylation near the TSS and near-complete loss of *SERPINH1* mRNA expression in differentiated MKs (**Figures 6G** and **S6D**). Although collagen-induced aggregation was modest in our assay, CDPs generated from *SERPINH1*-silenced cells exhibited a significantly reduced response relative to controls (**Figures 6H** and **S6E**).

## Discussion

Advances in HSC-based therapies have enabled one-time curative treatment of hematologic malignancies and inherited disorders. However, both the development of new engineering modalities and the application of HSC engineering to a broader range of diseases remain important opportunities. Compared with genome editing, epigenome editing modulates gene expression without altering the underlying DNA sequence and retains the capacity for reversibility, offering a potentially safer therapeutic modality. Within this class of approaches, DNA methylation-based silencers can achieve durable repression through deposition of heritable methylation marks following transient delivery.^13,14^ Such strategies have been successfully applied to silence disease-causing genes in post-mitotic tissues such as the brain, and in the liver, where cells undergo limited proliferation.^15–17^ More recently, durable epigenetic silencing has also been demonstrated in proliferating primary human T cells in proof-of-principle cell-engineering studies.^18,19^ However, the understudied HSCs represent a particularly demanding context, as methylation and silencing must be faithfully propagated through extensive cell division that both sustains self-renewal and generates differentiated hematopoietic progeny. Additionally, unlike prior applications in which the edited cells themselves constitute the relevant therapeutic target, HSC-based approaches require manipulation of a stem cell population to ultimately influence the function of downstream differentiated effector progeny. While a prior study demonstrated the feasibility of targeted DNA methylation editing in human HSPCs^20^, important questions remained regarding the robustness and durability of silencing, its reversibility and generalizability, as well as the broader translational potential of engineered epigenetic states in human hematopoiesis. We reasoned that platelet function would provide a compelling context in which to address these questions, as platelets are continuously derived from HSCs and play a central role in thrombosis. Durable modulation of platelet function through HSC epigenome editing could therefore offer a potential strategy for thrombosis prevention. Here, by targeting the platelet receptor integrin β3 (*ITGB3*), a central mediator of platelet aggregation, we show that its silencing by CHARM in human HSPCs can be durably inherited through HSC self-renewal and megakaryocytic differentiation, with functional consequences for platelet aggregation.

From a safety perspective, CHARM-mediated editing of *ITGB3* was highly specific. Methylome and transcriptome profiling revealed minimal off-target effects on DNA methylation and gene expression. At the targeted locus, we observed localized transcriptional changes, including notable up-regulation of the neighboring gene *MYL4*, likely due to local chromatin remodeling associated with perturbation of the *ITGB3* locus during MK differentiation. However, this on-target effect did not elicit secondary alterations in global transcriptional programs. Consistent with this specificity, CHARM editing did not affect HSC viability, apoptosis, short-or long-term HSC maintenance, colony-forming potential *in vitro*, or engraftment and multilineage reconstitution *in vivo. MYL4* encodes a member of the myosin light chain family and could, in principle, influence cytoskeletal or contractile properties. However, we did not observe broader dysregulation of myosin-associated or cytoskeletal gene programs, nor evidence that MYL4 upregulation impacted HSCs in a measurable manner. While more comprehensive characterization of the functional consequences of MYL4 upregulation will be important in future studies, the major phenotypes observed here were consistent with the expected effects of specific *ITGB3* silencing. Importantly, CHARM-mediated silencing is reversible through targeted demethylation, highlighting a key control feature of epigenome editing in human HSCs. While reversal of CD61 expression was partial under the conditions tested, potentially reflecting reduced editing efficiency following extended *in vitro* culture or less optimized demethylation kinetics, approximately 60% of cells reverted to a CD61^+^ state, demonstrating substantial restoration of gene expression.

Of note, we used *ITGB3* as a proof-of-principle target based on its CpG-rich promoter, well-defined role in platelet aggregation, favorable safety profile inferred from human genetics, and clinical precedent demonstrating that inhibition of αIIbβ3 reduces thrombotic events.^34,40,41,55^ Extending beyond this single target, we demonstrated the broader applicability of CHARM-mediated silencing to genes acting at different steps of thrombus formation and thus with distinct effectiveness-risk profiles. Target selection was also guided by the requirement for CpG-rich regulatory regions with low baseline methylation, and accordingly we focused on genetically inspired antiplatelet targets with annotated promoter-associated CpG islands. Future studies will be required to define the extent to which alternative regulatory architectures, including distal enhancers, can be leveraged to expand the range of therapeutically actionable targets. Identifying targets that provide the most favorable balance between antithrombotic efficacy and bleeding liability will require future studies integrating human genetics, clinical evidence, and detailed functional characterization.

Together, our study establishes durable and reversible epigenome editing of human HSCs as a platform for long-term modulation of platelet function, supporting its future investigation for thrombosis prevention. More broadly, it defines HSC-based epigenome editing as a general approach for sustained and controllable regulation of hematopoietic gene expression across diseases. These findings provide a foundation for future efforts to harness programmable epigenetic editing in HSCs for therapeutic benefit. While this study did not focus on delivery, we envision that the engineering of HSCs could be applied *ex vivo*, as current therapies are delivered, or *in vivo*, as emerging approaches for HSC delivery such as lipid nanoparticles and virus-like particles continue to rapidly advance.^56–58^

### Limitations of the study

While the NBSGW xenotransplantation model is optimized for supporting long-term human hematopoiesis, human megakaryopoiesis and platelet production only occur at low levels in this system, and circulating human platelets are extremely rare. As a result, *in vivo* functional assessment of platelet activity and thrombus formation following epigenome editing was not feasible in this model. In addition, because *AAVS1*-targeted CHARM, rather than unedited HSPCs, were used as controls throughout this study, potential target-independent effects of CHARM cannot be definitively excluded. Finally, although *ITGB3* served as a useful proof-of-principle target, this study did not define the degree of gene modulation required to achieve antithrombotic benefit, while preserving normal hemostasis. Future studies will be needed to determine the therapeutic window associated with different levels of target suppression and to establish the degree of reversal required for functional rescue.

## Acknowledgements

We thank members of the Sankaran lab and other colleagues for valuable advice and guidance. This work was supported by the Howard Hughes Medical Institute (to V.G.S.), National Institutes of Health grants R01DK103794, R01DK101989, R01HL146500, R01CA265726, R01CA292941 (to V.G.S.), the Gates Foundation (to V.G.S.), the Mathers Foundation (to V.G.S.), the Edward P. Evans Foundation (to V.G.S.), Blood Cancer United (to V.G.S.), Alex’s Lemonade Stand Foundation (to V.G.S.), the Leona and Harry Helmsley Charitable Trust (to V.G.S.), and a gift in memory of Jan Ellen Paradise, MD to Boston Children’s Hospital (to V.G.S.). V.G.S. is an Investigator of the Howard Hughes Medical Institute.

## Author contributions

Conceptualization, T.Y. and V.G.S.; Methodology, T.Y., M.N.B., R.M., A.A.S., K.R.M., and V.G.S.; Investigation, T.Y., M.N.B, P.L., M.A., L.d.V., C.G., E.V., I.A.J., M.S.T., S.D.S., L.W., A.C., L.M., G.A., and R.M.; Computational analyses, T.Y., W.X., A.J.L., and M.P.; Resources, M.N.B., A.A.S., J.S.W., and K.R.M.; Funding acquisition and Supervision, V.G.S.; Writing, T.Y. and V.G.S. with input from all authors.

## Declaration of Interests

V.G.S. is an advisor to Ensoma, Cellarity, and Beam Therapeutics, unrelated to this work. J.S.W. declares outside interest in 5 AM Ventures, Amgen, Chroma Medicine, KSQ Therapeutics, Maze Therapeutics, Tenaya Therapeutics, Tessera Therapeutics, Third Rock Ventures, and Xaira Therapeutics. There are no other relevant declarations to report.

## Resource availability

Plasmids generated in this study have been deposited with Addgene and will be publicly available at the time of publication. All data and code will be released at the time of publication.

## Materials and Methods

### Primary cell culture

All experiments were conducted in accordance with institutional guidelines and approval at Boston Children’s Hospital.

CD34^+^ HSPCs isolated from G-CSF-mobilized peripheral blood of adult donors (both sexes, 26-46 years of age) were purchased from the Fred Hutchinson Cancer Research Center. Cryopreserved cells were thawed rapidly in a 37 °C water bath and gradually diluted with thawing medium (DPBS + 1% FBS). Cells were then cultured at 37 °C in StemSpan Serum-Free Expansion Medium (SFEM) II (STEMCELL Technologies) supplemented with L-glutamine (2 mM), penicillin-streptomycin (100 U/mL), StemSpan CC100 (1×; STEMCELL Technologies), thrombopoietin (TPO) (50 ng/mL; PeproTech), and UM171 (35 nM; STEMCELL Technologies).

Cord blood was obtained from the Brigham and Women’s Hospital Center for Clinical Investigation. Donor sex and gestational age information for the cord blood units was unavailable. CD34^+^ cells were enriched using the EasySep Human CD34 Positive Selection Kit (STEMCELL Technologies) according to the manufacturer’s instructions and were either cryopreserved until use or directly cultured.

For chemically defined cytokine-free culture, cells were cultured in Iscove’s modified Dulbecco’s medium (IMDM; Gibco) supplemented with 1% insulin-transferrin-selenium-ethanolamine (Life Technologies), penicillin-streptomycin (100 U/mL), UM171 (70 nM), polyvinyl alcohol (1 mg/mL; Sigma-Aldrich), 740Y-P (1 µM; MedChemExpress), and butyzamide (0.1 µM; MedChemExpress). Cells were maintained at a density of 5 × 104 to 1 × 105 cells per mL in CellBIND plates (Corning).

For megakaryocytic differentiation, edited cells were cultured in StemSpan Serum-Free Expansion Medium (SFEM) II supplemented with penicillin-streptomycin (100 U/mL), human stem cell factor (SCF; 100 ng/mL), TPO (100 ng/mL), Flt-3 ligand (100 ng/mL; PeproTech), and IL-6 (100 ng/mL; PeproTech) for 6 days, followed by culture in StemSpan SFEM II supplemented with penicillin-streptomycin (100 U/mL) and TPO (50 ng/mL) for an additional 6 days. For proplatelet formation experiments, heparin (25 U/mL) was added to the latter differentiation medium. In experiments generating CDPs, SR1 (1 µM; MedChemExpress) and human low-density lipoprotein (LDL; 20 µg/mL; Sigma-Aldrich) were added to both media. Following the 12-day differentiation, cultures were maintained for an additional 3 days under orbital shaking with aspirin (200 µM; Cayman Chemical Company) to promote CDP production.

For erythroid differentiation, a previously described three-phase culture system was used.^59^ The base erythroid medium consisted of IMDM supplemented with 2% human AB plasma (SeraCare), 3% human AB serum (Thermo Fisher Scientific), heparin (3 U/mL), insulin (10 µg/mL), holo-transferrin (200 µg/mL), and penicillin-streptomycin (100 U/mL). During days 1-7, the base medium was further supplemented with erythropoietin (EPO; 3 U/mL), human SCF (10 ng/mL), and IL-3 (1 ng/mL). From days 8-12, cultures were maintained in base medium supplemented with EPO (3 U/mL) and human SCF (10 ng/mL). From days 13-20, cells were cultured in base medium supplemented with EPO (3 U/mL) and holo-transferrin at a final concentration of 1 mg/mL.

### Mouse models

All mouse procedures were conducted in accordance with protocols approved by the Institutional Animal Care and Use Committee at Boston Children’s Hospital.

CD34^+^ cells enriched from cord blood were nucleofected with CHARM mRNA together with sgRNA targeting either the *AAVS1* locus (control) or the *ITGB3* promoter. Two days after nucleofection, 2 × 105 CD34^+^ HSPCs were intravenously injected via the tail vein into eight-to ten-week-old male and female NBSGW mice. Sixteen weeks after transplantation and two days after administration of clodronate liposomes, mice were euthanized by CO_2_ inhalation, and total bone marrow cells were harvested from the ilia, femurs, and tibias. A fraction (3%) of the bone marrow was reserved for erythroid staining, while the remaining cells were subjected to red blood cell lysis followed by engraftment flow cytometric analysis, CD34^+^ cell enrichment, and downstream analyses. Both male and female mice were used for all experiments.

### mRNA production

Coding regions of CRISPRi, CRISPRoff, CHARM, and TET1-dCas9 were cloned into the PEmaxNG-_2star_IVT (Addgene) backbone, which contains sequences of a T7 promoter and optimized 5′ and 3′ untranslated regions (UTRs). Full-length DNA templates were generated by PCR amplification using a reverse primer containing a 5′ poly(T) tract to add a poly(A) tail. PCR products were purified using the Monarch PCR & DNA Cleanup Kit (NEB), and *in vitro* transcription was performed following the Standard-Yield Protocol associated with CleanCap Reagent M6 (TriLink), with N1-methylpseudouridine-5′-triphosphate substituted for UTP and reactions incubated at 37 °C for 3 h. Template DNA was removed by treatment with DNase I in the presence of 10× DNase buffer (NEB) at 37 °C for an additional 15 minutes. RNA was purified using the Monarch Spin RNA Cleanup Kit (NEB), quantified by spectrophotometry, and stored at −80 °C until use.

### Editing of CD34^+^ HSPCs

sgRNAs targeting sites within approximately 200 bp of the TSS were designed and prioritized based on predicted on-target activity and off-target specificity.^60,61^ For each target, two to three candidate sgRNAs within the desired window were typically evaluated, and the sgRNA producing the most robust target-gene silencing was selected. Electroporation of human CD34^+^ HSPCs was performed using the Lonza 4D-Nucleofector system with the P3 Primary Cell 4D-Nucleofector kit (20 µL format). Briefly, cells were harvested, washed twice with DPBS, and resuspended in P3 Primary Cell Nucleofector Solution supplemented with Supplement 1. Cell suspensions were mixed with 100 pmol chemically modified sgRNAs (Synthego) and 2 µg of purified mRNA, transferred to cuvettes, and nucleofected using program DS130.

### Cell staining and flow cytometric analysis

For surface staining, cells were harvested and washed with FACS buffer (DPBS + 2% FBS + 2 mM EDTA), then resuspended in a panel of fluorophore-conjugated antibodies diluted in FACS buffer. Samples were stained at room temperature for 30 min protected from light, washed, resuspended in FACS buffer, and analyzed on a BD LSRFortessa or Cytek Aurora flow cytometer. Data were analyzed using FlowJo.

For division tracing, one day after nucleofection, cells were labeled with CellTrace CFSE (Thermo Fisher) according to the manufacturer’s instructions. Briefly, cells were harvested, washed twice with DPBS, and resuspended in 0.5 µM CFSE diluted in DPBS. After incubation at 37 °C for 10 minutes protected from light, labeling was quenched by the addition of five volumes of FACS buffer. Cells were then washed twice with DPBS and returned to culture. CFSE fluorescence was assessed by flow cytometric analysis on a Cytek Aurora flow cytometer at day 0 and day 5 after labeling. Data were analyzed using FlowJo.

For apoptosis/viability analysis, cells were harvested and washed once with Annexin V binding buffer (BioLegend), then resuspended in Annexin V binding buffer containing Annexin V and DAPI. Samples were incubated for 15 min at room temperature protected from light, and analyzed on a Cytek Aurora flow cytometer. Data were analyzed using FlowJo.

### Colony formation assay

Colony formation assays were performed using MethoCult H4034 Optimum (STEMCELL Technologies) according to the manufacturer’s instructions. Specifically, two days after electroporation, cells were counted, mixed with MethoCult medium, and evenly dispensed into SmartDish (STEMCELL Technologies), with 500 cells plated per well. Colonies were imaged after 12-15 days and harvested by washing with cold DPBS. For serial replating assays, secondary and tertiary colony formation was performed under the same conditions, except that 50,000 and 300,000 cells were used for both secondary and tertiary platings, respectively.

### Immunofluorescence (IF) imaging of megakaryocyte and proplatelets

At the end of megakaryocytic differentiation, cells were harvested and size-selected using a 20 µm cell strainer. The retained fraction was fixed with 4% paraformaldehyde diluted in DPBS at room temperature for 30 min, washed once with DPBS, and blocked in 3% BSA diluted in DPBS at room temperature for 30 min. Cells were then stained with the indicated antibodies overnight at 4 °C. After washing, cells were resuspended in DPBS and imaged.

For proplatelet formation and imaging, Lab-Tek chamber slides were coated with anti-CD31 antibody at 37 °C for 1 h and blocked with 3% BSA diluted in PBS. Cells were harvested, size-selected using a 20 µm cell strainer, and seeded onto the coated chambers. Cells were incubated overnight at 37 °C to allow proplatelet formation. Culture medium was then gently removed, and cells were fixed with 4% paraformaldehyde diluted in DPBS at room temperature for 30 min, washed once with DPBS, and blocked with 3% BSA diluted in DPBS at room temperature for 30 min. Cells were incubated with the indicated antibodies overnight at 4 °C. After removal of the antibody solution, cells were maintained in DPBS and imaged.

Fluorescence Z-stacks were processed in Fiji and displayed as maximum intensity projections; sg*AAVS1* and sg*ITGB3* samples were imaged using identical acquisition settings, and brightness and contrast were adjusted for visualization.

### Targeted bisulfite sequencing

Genomic DNA (gDNA) was extracted from cultured or sorted cells using Quick-DNA Microprep Plus Kit or Quick-DNA/RNA Microprep Plus Kit (Zymo). For each sample, 200-1000 ng of gDNA was subjected to bisulfite conversion using the EZ DNA Methylation-Lightning Kit (Zymo) following the manufacturer’s instructions. Targeted regions were PCR-amplified using primers designed with Bisulfite Primer Seeker (Zymo) and Q5U Hot Start High-Fidelity DNA Polymerase (NEB). Amplicons were sequenced, and raw FASTQ files were filtered by read length and analyzed using QUMA to quantify CpG methylation status. Methylation at each CpG site was calculated as the percentage of reads scored as methylated among all informative reads covering that position.

### qPCR

RNA was isolated from cultured or flow-sorted cells using the Quick-RNA Microprep Plus Kit (Zymo). Equal amounts of RNA (up to 1 µg) were subjected to reverse transcription using PrimeScript RT Master Mix (TaKaRa). Quantitative PCR was performed using primers specifically targeting the indicated genes and SYBR Green Universal Master Mix (Thermo Fisher Scientific). Relative gene expression levels were normalized to *GAPDH* and calculated using the ΔΔCt method, with control samples serving as the reference.

### RNA sequencing

For MKs, RNA was extracted from CHARM-edited, differentiated, and sorted MKs using the Total RNA Purification Micro Kit (Norgen Biotek) according to the manufacturer’s instructions. Samples were processed and sequenced using Genewiz RNA-seq service. For HSPCs, samples were processed and sequenced using Plasmidsaurus RNA-seq service. All RNA-seq data were aligned to the human reference genome (hg38) using STAR^62^ with the parameter --runMode alignReads --chimSegmentMin 20. The resulting alignments were sorted, and unmapped reads were removed using SAMtools^63^ with the option -F 1804. Gene-level read counts were then quantified using featureCounts^64^ based on the human reference gene annotation. Raw read counts were normalized to Transcripts Per Million (TPM) to account for both sequencing depth and gene length. For each gene, read counts were first adjusted by gene length to obtain expression values in reads per kilobase. These values were then scaled by the total expression of all genes in the same sample and multiplied by one million to generate TPM values, ensuring that the sum of TPM values within each sample equals one million and allowing direct comparison of gene expression levels across samples.

### Whole-genome bisulfite sequencing

gDNA was extracted from CHARM-edited HSPCs using Quick-DNA Microprep Plus Kit (ZYMO) 4 days post electroporation. Samples were processed and sequenced using Genewiz WGBS service. Raw WGBS-seq FASTQ files were aligned to the reference genome using Bismark^65^ (https://felixkrueger.github.io/Bismark/bismark/alignment/) with Bowtie2^66^ in directional mode and default parameters (--score_min L,0,-0.2, paired-end minimum/maximum insert size: -I 0, -X 500). CpG methylation levels were extracted from Bismark coverage files and imported into the methylKit^67^ (https://github.com/al2na/methylKit) R package for downstream analysis. To identify differentially methylated tiles, the tileMethylCounts function was applied with a window size of 1000 bp and a step size of 100 bp. Differentially methylated regions (DMRs) were detected using the calculateDiffMeth function with overdispersion = “MN” and adjust = “BH” for multiple testing correction. DMRs were ranked based on absolute methylation percentage differences, and statistical significance was defined as a false discovery rate (FDR) < 0.01 with an absolute methylation difference ≥25%. Results were visualized using Manhattan plots displaying the −log10-transformed p-values of individual tiling windows. For locus-level visualization on UCSC genome browser, base-resolution methylation information was extracted from Bismark BedGraph files and converted into a compatible format. Methylation levels were displayed as bar plots, where hypermethylated regions (50%–100% methylation) were shown in red on a scale from 0.5 to 1, and hypomethylated regions (0%–50% methylation) were shown in blue on a scale from −1 to −0.5.

### CDP activation and fibrinogen binding assay

CDPs generated as described above were harvested following the addition of prostaglandin I_2_ (PGI_2_; Cayman Chemical Company) and apyrase (Sigma-Aldrich). Samples were centrifuged at 300 g for 5 minutes to remove intact cells, and CDPs were subsequently pelleted by centrifugation at 1,000 g for 10 minutes (low brake). Agonists were added as indicated (ADP, 200 µM; TRAP, 200 µM; U46619, 5 µM) together with antibodies against CD42b, Annexin V, P-selectin, and PAC-1 for activation assays or fibrinogen Alexa Fluor 647 conjugate for fibrinogen-binding assays, followed by incubation for 15 min at room temperature or 37°C, respectively. Samples were then fixed with 0.5% paraformaldehyde and analyzed on a Cytek Aurora flow cytometer. Data were analyzed using FlowJo software.

### Platelet aggregation assay

CDPs were harvested as described above. Whole blood was collected into BD Vacutainer Citrate Tubes and centrifuged at 210 g for 15 minutes (low brake). Platelet-rich plasma (PRP) was collected, and modified Tyrode’s buffer (MTB) supplemented with PGI_2_ and apyrase was added. Platelets were then pelleted by centrifugation at 1,000 g for 10 min (low brake).

CDPs and platelets were stained with CD31-Pacific Blue and PE, respectively, for 15 minutes at room temperature and washed with MTB. Cells were resuspended in MTB supplemented with CaCl_2_ (2 mM), human fibrinogen (0.1 mg/mL), CPD plasma (25% vol/vol), and 5 × 10^6^ platelets were combined with an equal number of CDPs from different editing. Agonists were added as indicated (ADP 10µM, TRAP 10 µM, U46619 0.25 µM, Ristocetin 1.5 mg/mL, Collagen 10 µg/mL), and samples were incubated at 37 °C with shaking at 800 rpm in a thermomixer for 5 minutes. Reactions were fixed by the addition of five volumes of 0.5% paraformaldehyde and analyzed on a Cytek Aurora flow cytometer. Data were analyzed using FlowJo.

### Quantification and statistical analysis

For experiments using human primary CD34^+^ HSPCs, data were obtained from at least two independent donors unless otherwise indicated. For xenotransplantation studies, seven recipient mice per group, including both male and female animals, were analyzed. For comparisons between two groups, statistical significance was assessed using a two-tailed unpaired t-test with Welch’s correction. For experiments involving more than two groups or multiple experimental conditions, one-way or two-way ANOVA was performed, followed by post hoc multiple-comparison correction. All data are presented as mean ± SD, significance is indicated as \**P* < 0.05, \*\**P* < 0.01, \*\*\**P* < 0.001, or n.s. not significant.

## Supplemental information

Table S1. Primer and sgRNA sequences used in the study

Table S2. Differentially methylated regions identified in HSPCs edited with CHARM targeting the *ITGB3* promoter or *AAVS1* as a control

Table S3. RNA-seq analyses in MKs and HSPCs edited with CHARM targeting the *ITGB3* promoter or *AAVS1* as a control

Table S4. Bisulfite sequencing results

## Supplemental Figures

**Figure S1.**
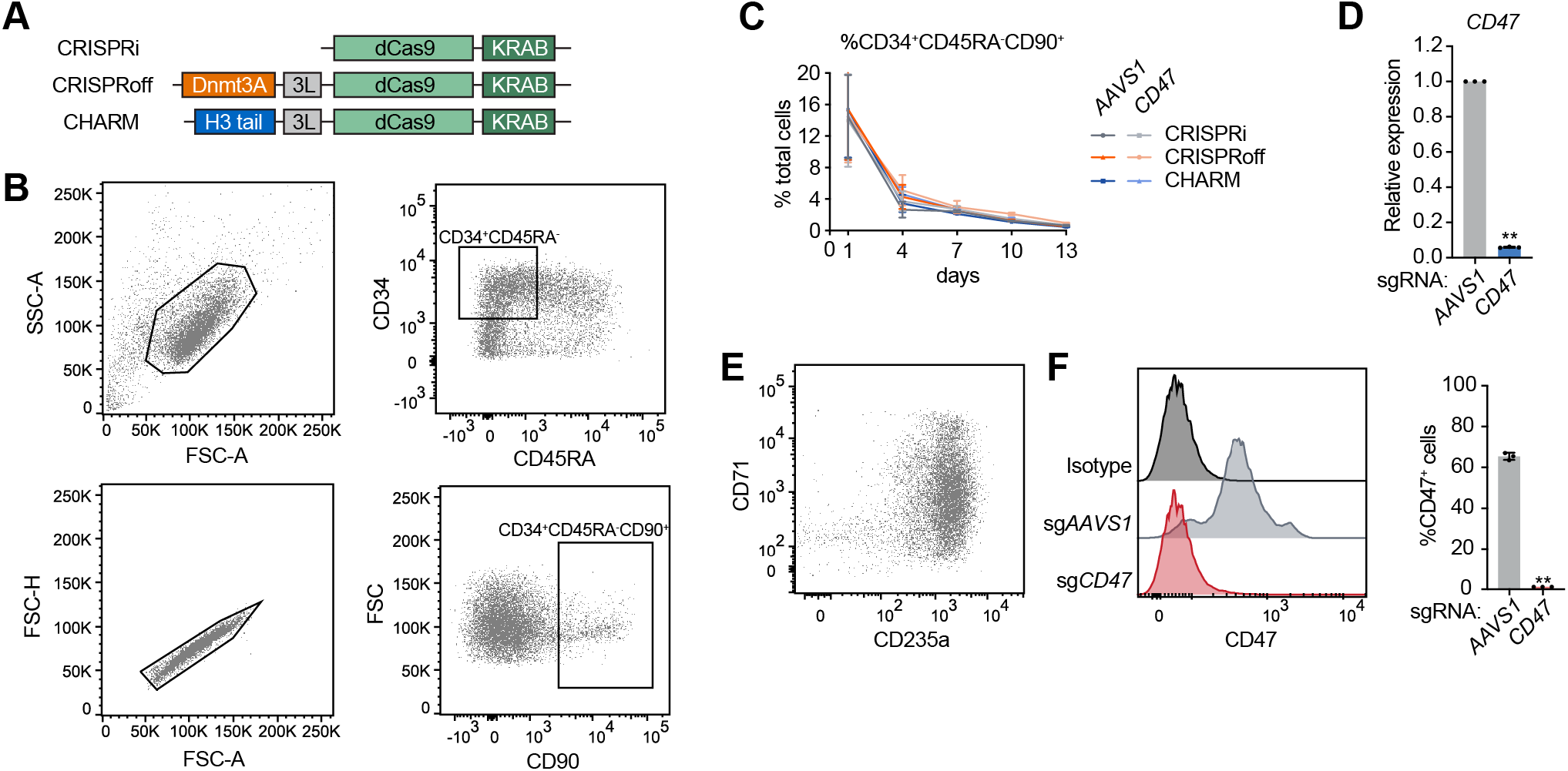
CHARM enables durable epigenetic silencing in human HSPCs. **(A)** Schematic representation of CRISPRi, CRISPRoff, and CHARM constructs used in experiments. **(B)** Representative flow cytometry gating strategy used to define CD34^+^CD45RA^−^CD90^+^ cells within the HSPC culture. **(C)** Quantification of the percentage of CD34^+^CD45RA^−^CD90^+^ cells over time in HSPC cultures following editing with CRISPRi, CRISPRoff, or CHARM targeting the *CD47* promoter or *AAVS1* as a control. **(D)** Quantification of *CD47* mRNA expression measured by RT-qPCR in HSPCs 13 days after editing with CHARM targeting the *CD47* promoter or *AAVS1* as a control. **(E)** Representative flow cytometry plots showing erythroid differentiation profiles based on CD71 and CD235a expression following differentiation of CHARM-edited HSPCs. **(F)** Left, representative flow cytometry histograms of CD47 surface expression in erythroid cells differentiated from HSPCs edited with CHARM targeting the *CD47* promoter or *AAVS1* as a control. Right, quantification of the percentage of CD47^+^ cells. All data are presented as mean ± SD, significance is indicated as \**P* < 0.05, \*\**P* < 0.01, or n.s. not significant.

**Figure S2.**
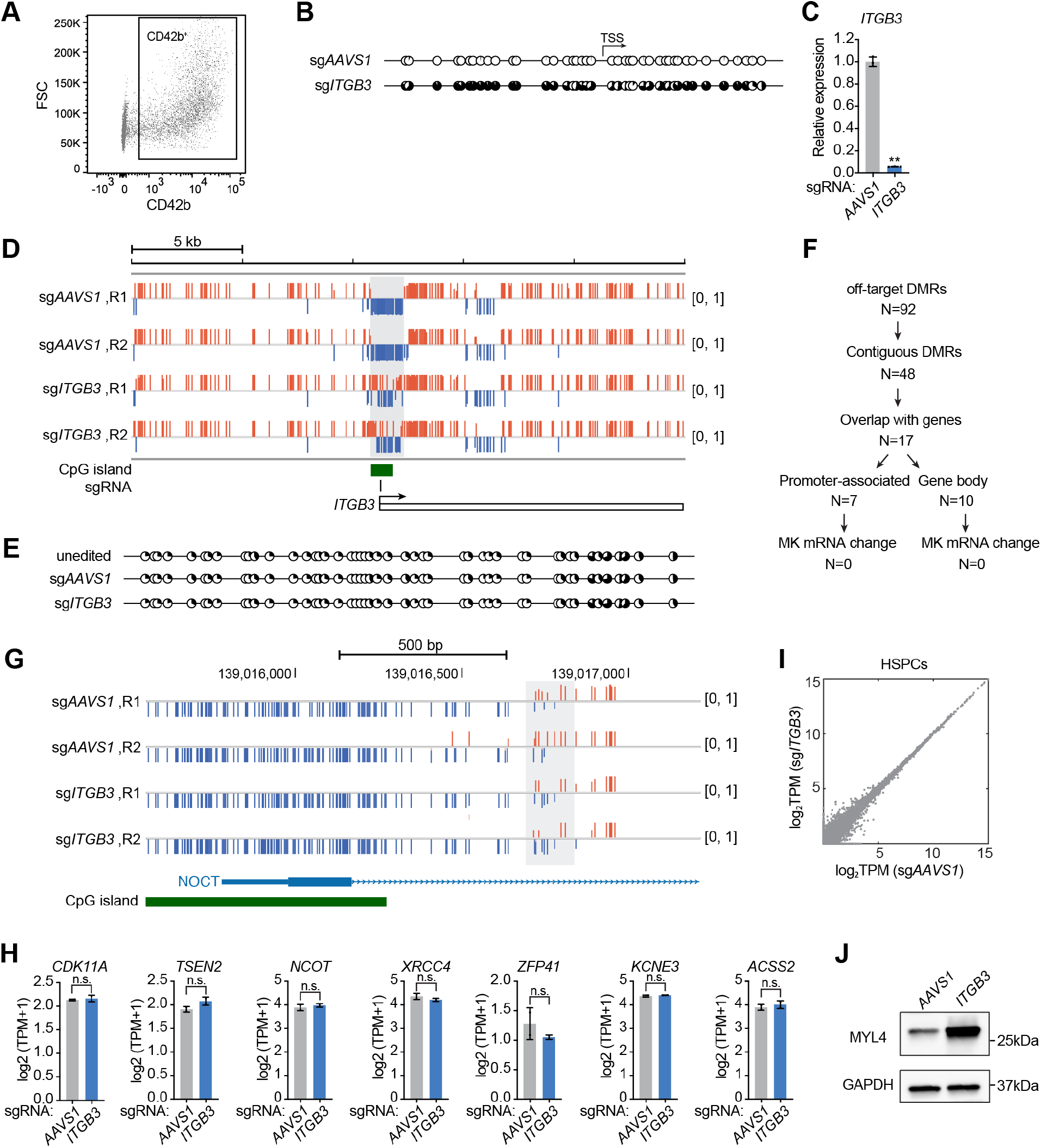
Robust and specific silencing of *ITGB3* by CHARM persists through megakaryocytic differentiation. **(A)** Representative flow cytometry gating strategy used to define CD42b^+^ mature megakaryocytes following differentiation of CHARM-edited HSPCs. **(B)** Representative targeted bisulfite sequencing displaying CpG methylation patterns near the *ITGB3* TSS in megakaryocytes differentiated from HSPCs edited with CHARM targeting the *ITGB3* promoter or *AAVS1* as a control. Each circle represents an individual CpG site; filled circles indicate methylated CpGs. *P* < 0.0001. **(C)** Quantification of *ITGB3* mRNA expression measured by RT-qPCR in differentiated megakaryocytes following CHARM targeting of the *ITGB3* promoter or *AAVS1* as a control. **(D)** Genome browser view comparing CpG methylation across a genomic window spanning the *ITGB3* locus in HSPCs edited with CHARM targeting the *ITGB3* promoter or *AAVS1* as a control. Red and blue marks denote CpGs with high (β-value > 0.5) or low (β-value < 0.5) DNA methylation, respectively. The annotated CpG island is shown in green, and the sgRNA target site is indicated. **(E)** Targeted bisulfite sequencing displaying CpG methylation patterns in the chromosome 18q21.32 region identified by WGBS in unedited HSPCs and HSPCs edited with CHARM targeting *AAVS1* or *ITGB3*. Each circle represents an individual CpG site; filled circles indicate methylated CpGs. **(F)** Hierarchical annotation of off-target differentially methylated regions (DMRs) identified by WGBS in CHARM-edited HSPCs. The number of DMRs at each step is indicated. (**G)** Genome browser view comparing CpG methylation across a representative off-target locus (*NOCT*) in HSPCs edited with CHARM targeting the *ITGB3* promoter or *AAVS1* as a control. Tracks show two replicates per condition. Red and blue marks denote CpGs with high (β-value > 0.5) or low (β-value < 0.5) DNA methylation, respectively. The annotated CpG island is shown in green. The grey shaded region highlights the area differentially methylated. **(H)** Expression of genes associated with promoter-overlapping differentially methylated regions identified by WGBS in HSPCs edited with CHARM targeting the *ITGB3* promoter or *AAVS1* as a control, measured by RNA sequencing. Transcript abundance is shown as log2(TPM + 1). **(I)** Scatter plot showing transcript abundance (log2(TPM + 1)) for all genes in HSPCs edited with CHARM targeting the *ITGB3* promoter or *AAVS1* as a control. **(J)** Immunoblot analysis of MYL4 protein expression in differentiated megakaryocytes following CHARM targeting of the *ITGB3* promoter or *AAVS1* as a control. GAPDH serves as a loading control. All data are presented as mean ± SD, significance is indicated as \**P* < 0.05, \*\**P* < 0.01, or n.s. not significant.

**Figure S3.**
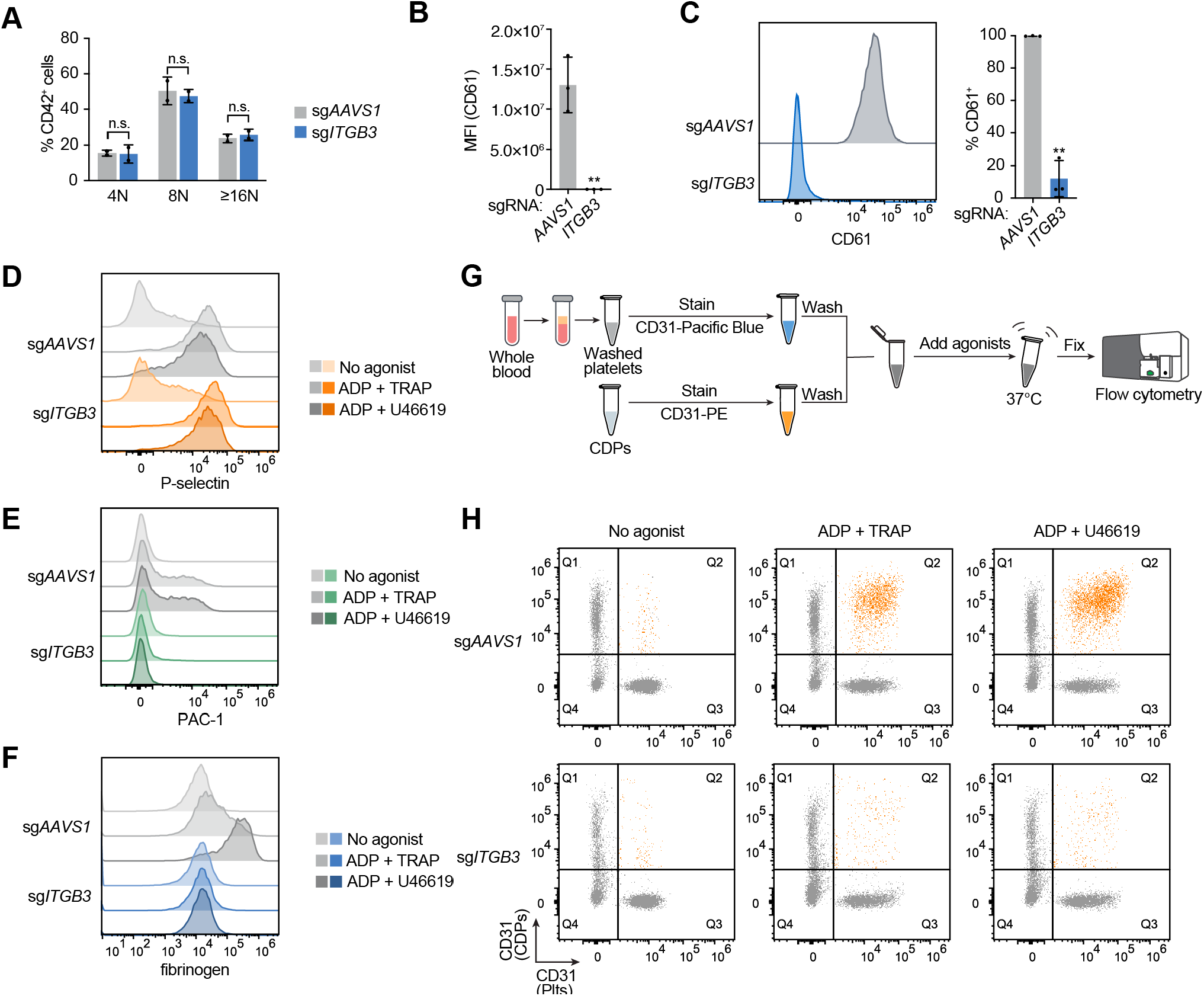
Silencing of *ITGB3* by CHARM persists through megakaryocytic differentiation and impairs αIIbβ3-dependent platelet function. **(A)** Quantification of megakaryocyte ploidy distribution following differentiation of HSPCs edited with CHARM targeting the *ITGB3* promoter or *AAVS1* as a control. Data are shown as the percentage of CD42^+^ cells within each ploidy class (n = 2 donors). **(B)** Quantification of CD61 MFI measured by IF imaging in 20-30 individual megakaryocytes identified by CD42d staining per condition. **(C)** Left, representative flow cytometry histograms of CD61 (*ITGB3*) surface expression in CDPs derived from HSPCs edited with CHARM targeting the *ITGB3* promoter or *AAVS1* as a control. Right, quantification of the percentage of CD61^+^ cells among CDPs. **(D)** Representative flow cytometry histograms of P-selectin induction in CDPs generated from HSPCs edited with CHARM targeting the *ITGB3* promoter or *AAVS1* as a control under no-agonist conditions or following stimulation with ADP + TRAP or ADP + U46619. **(E)** Representative flow cytometry histograms of PAC-1 binding in CDPs generated from HSPCs edited with CHARM targeting the *ITGB3* promoter or *AAVS1* as a control under no-agonist conditions or following stimulation with ADP + TRAP or ADP + U46619. **(F)** Representative flow cytometry histograms of fibrinogen binding in CDPs generated from HSPCs edited with CHARM targeting the *ITGB3* promoter or *AAVS1* as a control under no-agonist conditions or following stimulation with ADP + TRAP or ADP + U46619. **(G)** Schematic overview of the flow cytometry-based platelet aggregation assay used to assess the ability of CDPs to aggregate with donor platelets. **(H)** Representative flow cytometry plots assessing platelet-CDP aggregation under no agonist conditions or following stimulation with ADP + TRAP or ADP + U46619 for CDPs generated from HSPCs edited with CHARM targeting the *ITGB3* promoter or *AAVS1* as a control. All data are presented as mean ± SD, significance is indicated as \**P* < 0.05, \*\**P* < 0.01, or n.s. not significant.

**Figure S4.**
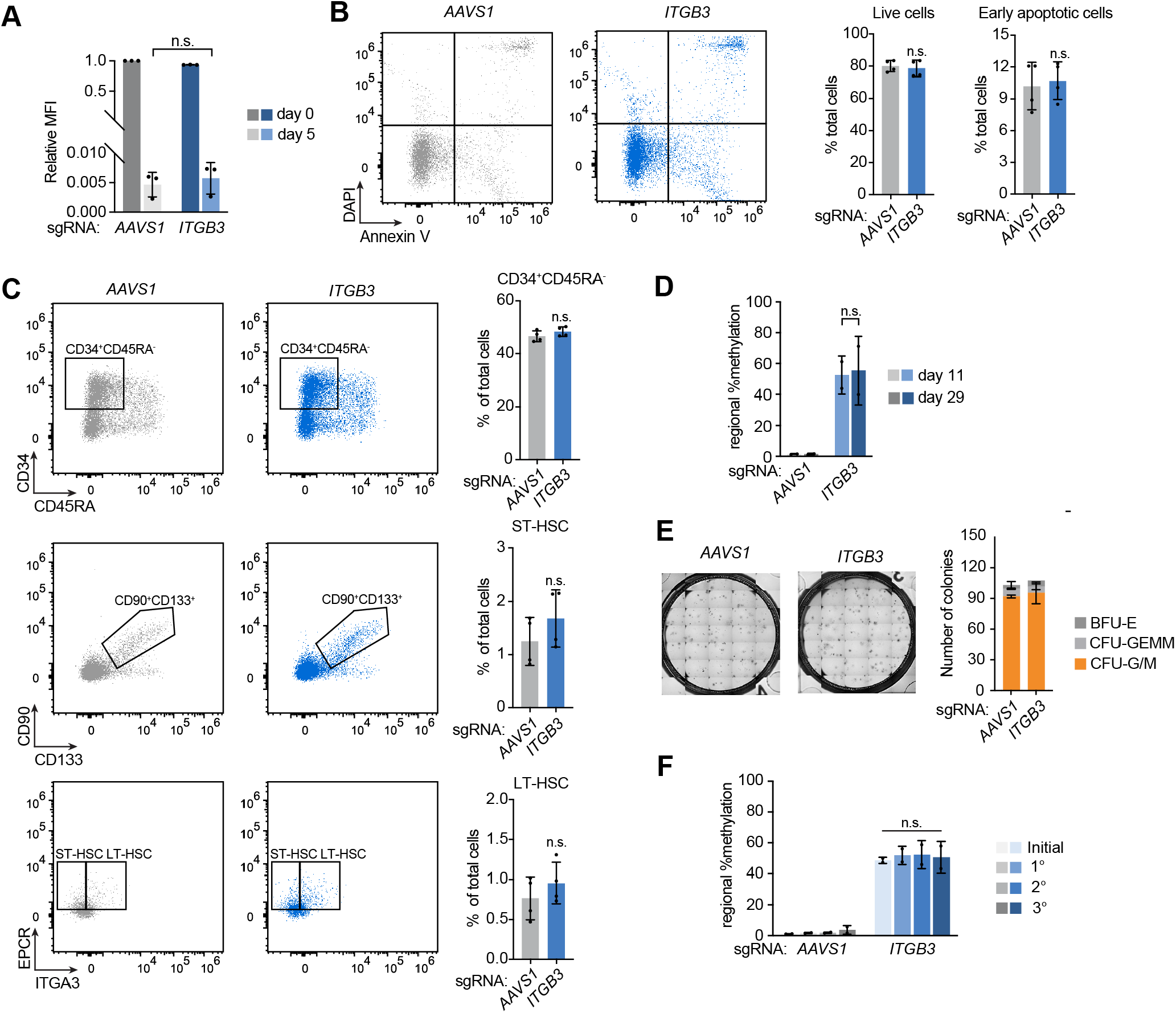
CHARM-mediated *ITGB3* silencing persists through HSC self-renewal. **(A)** Quantification of CFSE fluorescence intensity in HSPCs at day 0 and day 5 following CHARM targeting of the *ITGB3* promoter or *AAVS1* as a control, shown as normalized MFI. **(B)** Left, representative flow cytometry plots of Annexin V and DAPI staining in HSPCs edited with CHARM targeting *ITGB3* or *AAVS1* as a control. Right, quantification of live cells and early apoptotic cells. **(C)** Left, representative flow cytometry plots of CD34^+^CD45RA^−^, LT-HSC (CD34^+^CD45RA^−^CD90^+^CD133^+^EPCR^+^ITGA3^+^) and ST-HSC (CD34^+^CD45RA^−^CD90^+^CD133^+^EPCR^+^ITGA3^−^) in HSPCs edited with CHARM targeting *ITGB3* or *AAVS1* as a control. Right, quantification of the indicated populations as a percentage of total cells. **(D)** Quantification of regional DNA methylation near the *ITGB3* TSS in HSPCs at day 11 and day 29 following CHARM targeting of the *ITGB3* promoter or *AAVS1* as a control. **(E)** Top, quantification of colony-forming units (BFU-E, CFU-GEMM, and CFU-G/M) generated from HSPCs edited with CHARM targeting the *ITGB3* or *AAVS1* as a control. Bottom, representative images of methylcellulose cultures. **(F)** Quantification of regional DNA methylation near the *ITGB3* TSS following CHARM targeting of the *ITGB3* promoter or *AAVS1* as a control in the initial edited HSPC population and in cells recovered from primary (1°), secondary (2°), and tertiary (3°) CFU replatings. All data are presented as mean ± SD, significance is indicated as \**P* < 0.05, \*\**P* < 0.01, or n.s. not significant.

**Figure S5.**
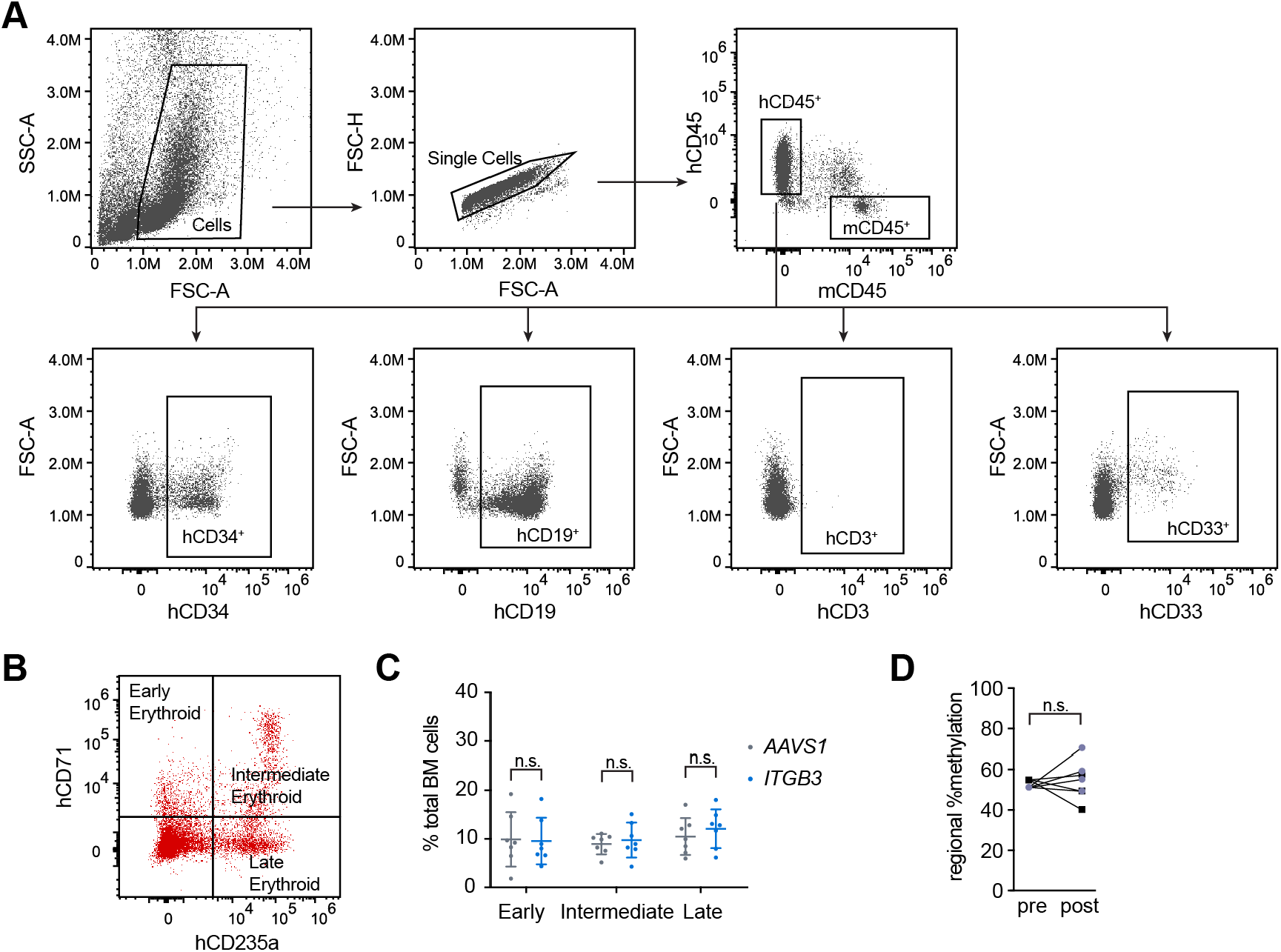
*ITGB3* silencing by CHARM is maintained following long-term engraftment in xenotransplanted mice. **(A)** Representative flow cytometry gating strategy used to identify human hematopoietic populations in the bone marrow of transplanted NBSGW mice, including hCD45^+^ cells and lineage-defined subsets (hCD34^+^, hCD19^+^, hCD3^+^, and hCD33^+^). **(B)** Representative flow cytometry profiles showing erythroid differentiation stages in the bone marrow of recipient mice, defined by hCD71 and hCD235a expression. **(C)** Quantification of early, intermediate, and late erythroid populations as a percentage of total bone marrow cells in mice transplanted with CHARM-edited HSPCs targeting the *ITGB3* promoter or *AAVS1* as a control. **(D)** Quantification of regional DNA methylation near the *ITGB3* TSS in cells before transplantation (pre) and in CD34^+^ cells recovered from recipient bone marrow 16 weeks after transplantation (post) following CHARM targeting of the *ITGB3* promoter. All data are presented as mean ± SD, significance is indicated as \**P* < 0.05, \*\**P* < 0.01, or n.s. not significant.

**Figure S6.**
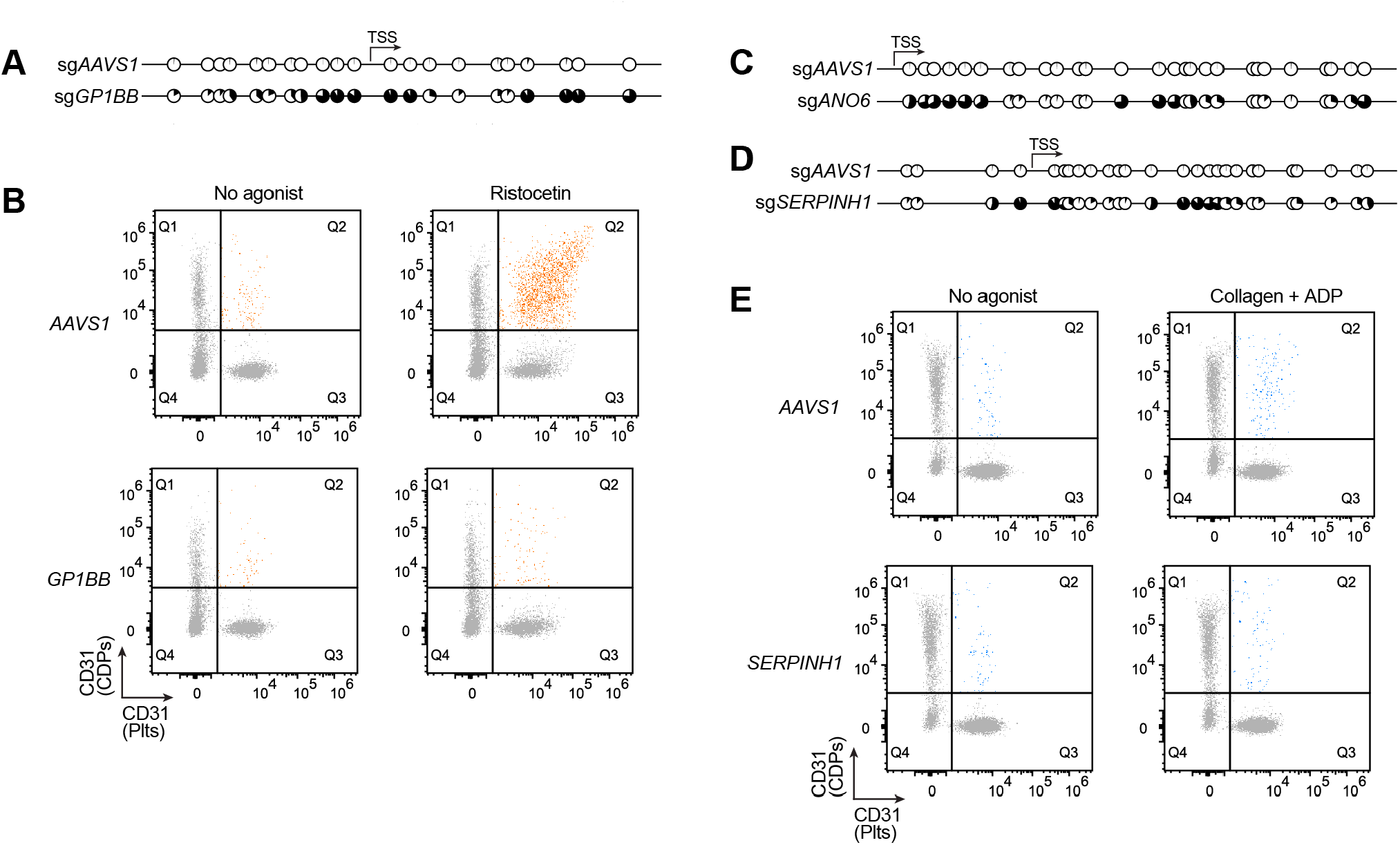
CHARM is broadly applicable for epigenetic silencing of multiple platelet genes in HSPCs. (**A)** Representative targeted bisulfite sequencing displaying CpG methylation patterns near the *GP1BB* TSS in megakaryocytes differentiated from HSPCs edited with CHARM targeting the *GP1BB* promoter or *AAVS1* as a control. Each circle represents an individual CpG site; filled circles indicate methylated CpGs. *P* < 0.0001. **(B)** Representative flow cytometry plots assessing platelet-CDP aggregation under no agonist conditions or following stimulation with ristocetin for CDPs generated from HSPCs edited with CHARM targeting the *GP1BB* promoter or *AAVS1* as a control. **(C–D)** Representative targeted bisulfite sequencing displaying CpG methylation patterns near the *ANO6* (C) and *SERPINH1* (D) TSSs in megakaryocytes differentiated from HSPCs edited with CHARM targeting the respective promoters or *AAVS1* as a control. Each circle represents an individual CpG site; filled circles indicate methylated CpGs. *P* < 0.0001 for both. **(E)** Representative flow cytometry plots assessing platelet-CDP aggregation under no agonist conditions or following stimulation with collagen + ADP for CDPs generated from HSPCs edited with CHARM targeting the *SERPINH1* promoter or *AAVS1* as a control.

## Notes

### Summary of Updates

This version has been revised to reflect the peer-reviewed manuscript accepted for publication.

